# Integrated Proteomics Identifies Neuritin (NRN1) as a Mediator of Cognitive Resilience to Alzheimer’s Disease

**DOI:** 10.1101/2022.06.15.496285

**Authors:** Cheyenne Hurst, Derian A. Pugh, Measho H. Abreha, Duc M. Duong, Eric B. Dammer, David A. Bennett, Jeremy H. Herskowitz, Nicholas T. Seyfried

## Abstract

We present an integrative proteomic strategy for the nomination and validation of proteins associated with cognitive resilience to Alzheimer’s disease (AD). Correlation network analysis across distinct stages of AD was used to prioritize protein modules linked to resilience. Neuritin (NRN1), a hub protein in a module associated with synaptic biology, was identified as a top candidate of resilience and selected for functional validation in cultured neurons. NRN1 provided dendritic spine resilience against amyloid-β (Aβ), and NRN1 blocked Aβ-induced neuronal hyperexcitability. The impact of exogenous NRN1 on the proteome of cultured neurons was assessed and integrated with the AD brain network. This revealed over-lapping synapse-related biology that linked NRN1-induced changes in cultured neurons with human pathways associated with AD resilience. Collectively, this highlights the utility of integrating the proteome from human brain and model systems to prioritize therapeutic targets that mediate resilience to AD.

## Introduction

As the aging population continues to expand, the public health burden of Alzheimer’s disease (AD) is projected to reach staggering numbers without the advent of effective disease altering therapies (1). AD is an irreversible neurodegenerative disease defined by its pathological hallmarks, amyloid-beta (Aβ) plaques and tau neurofibrillary tangles (NFTs) (2). Functional imaging and biomarker studies suggest AD pathological brain changes could initiate up to two decades before symptom onset, indicating a protracted prodromal disease phase ideal for early intervention (3). Importantly, many older individuals without dementia or mild cognitive impairment meet pathologic criteria for AD. Approximately one-third of individuals harbor high levels of AD and related disease pathology in their brains at autopsy but showed little to no signs of cognitive impairment in their lifetime (4, 5). These cognitively normal people with AD pathology are described as preclinical or asymptomatic AD (AsymAD) and appear to exhibit cognitive resilience to the clinical manifestations of AD dementia. One working hypothesis is that such individuals possess physiological resilience that confers the ability to maintain cognitive function despite the accumulation of AD-related pathologies (6–8). Identifying the specific mechanisms by which older individuals with AD pathology avoid dementia onset is one of the most pivotal, unanswered questions in the field.

Cognitive impairment in AD is the result of lost neuronal connectivity in brain regions critical to memory and other cognitive processes. For cognitive impairment to develop, there must be loss or dysfunction of the neural elements that subserve cognition, e.g., neurons, synapses and dendritic spines. Our work and that of others demonstrate preservation of neuron numbers and synaptic markers as well as enhanced dendritic spine remodeling in resilient cases (9–11). Together this implies that the ability to maintain cognitive function in an environment of AD pathology is linked to the preservation and maintenance of synapses or spines. These findings raise important questions: 1) what are the molecular pathways that drive preservation of synaptic connections and maintenance of cognitive abilities in resilient individuals? 2) How can we identify protein targets to exploit these mechanisms for therapies in at risk patients?

To address these gaps in knowledge, we implemented an integrative systems-level analysis of multi-region postmortem human brain proteomics derived from the Religious Order and Rush Memory and Aging Project (ROSMAP) to identify proteins and pathways significantly altered in resilient cases. ROSMAP is an information-rich longitudinal cohort-based study in which participants enroll without dementia, undergo annual cognitive and clinical assessments and donate their brains at death (12). Multiplex tandem mass tag mass spectrometry (TMT-MS)-based proteomic data was implemented for a correlation network analysis. Data from an independent brain proteome wide association study (PWAS) of cognitive trajectory was integrated with the brain network to robustly prioritize protein communities associated with cognitive resilience. This revealed proteins linked to synaptic biology and cellular energetics. Neuritin (NRN1) was prioritized as a hub that co-expressed with a community of proteins with high correlation to cognitive stability in life and is known for important roles in synaptic maturation and stability (13–15). To further validate our systems-level analysis, primary neuronal culture was used to evaluate neuroprotective mechanisms of NRN1. Primary neurons treated with NRN1 were protected against Aβ oligomer-induced retraction of dendritic spines and neuronal hyperexcitability. Targets engaged by NRN1 treatment also significantly overlapped with human modules of resilience. The current work establishes a pipeline for network-driven nomination of critical proteins and mechanisms influencing resilience and incorporates experimental validation to assess neuroprotective capacity and recapitulation of cellular phenotypes of resilience.

## Results

### Proteomic measurements align with neuropathological scores

Matched post-mortem brain tissue samples from Brodmann area 6 (BA6) and Brodmann area 37 (BA37) from 109 Religious Orders Study and Rush Memory and Aging Project (ROSMAP, n=218 samples total) cases were analyzed using tandem mass tag mass spectrometry (TMT-MS; **Fig. 1A**). BA6 is a frontal cortex area containing the premotor and supplementary motor cortices, important for roles in motor, language and memory functions (16). BA37 resides in the temporal cortex and contains the fusiform gyrus which has been linked to disrupted language and memory function in AD (17). Cases were classified as Control, asymptomatic Alzheimer’s disease (AsymAD) or Alzheimer’s disease (AD) based on semi-quantitative measures of amyloid (CERAD) and tau (Braak) deposition as well as cognitive function near time of death (18, 19). This classification strategy, similar in concept to the A/T/N framework, which stratifies cases based on presence or absence of amyloid, tau and neurodegeneration, allows distinction of cases with increased neuropathological burden but intact cognitive function (20). Protein expression data was adjusted for batch-effects, outlier removal and confounding effects of covariates (age, sex and post-mortem interval or PMI) for a final expression dataset of 7,787 proteins (**Fig. S1**).

**Figure 1.**
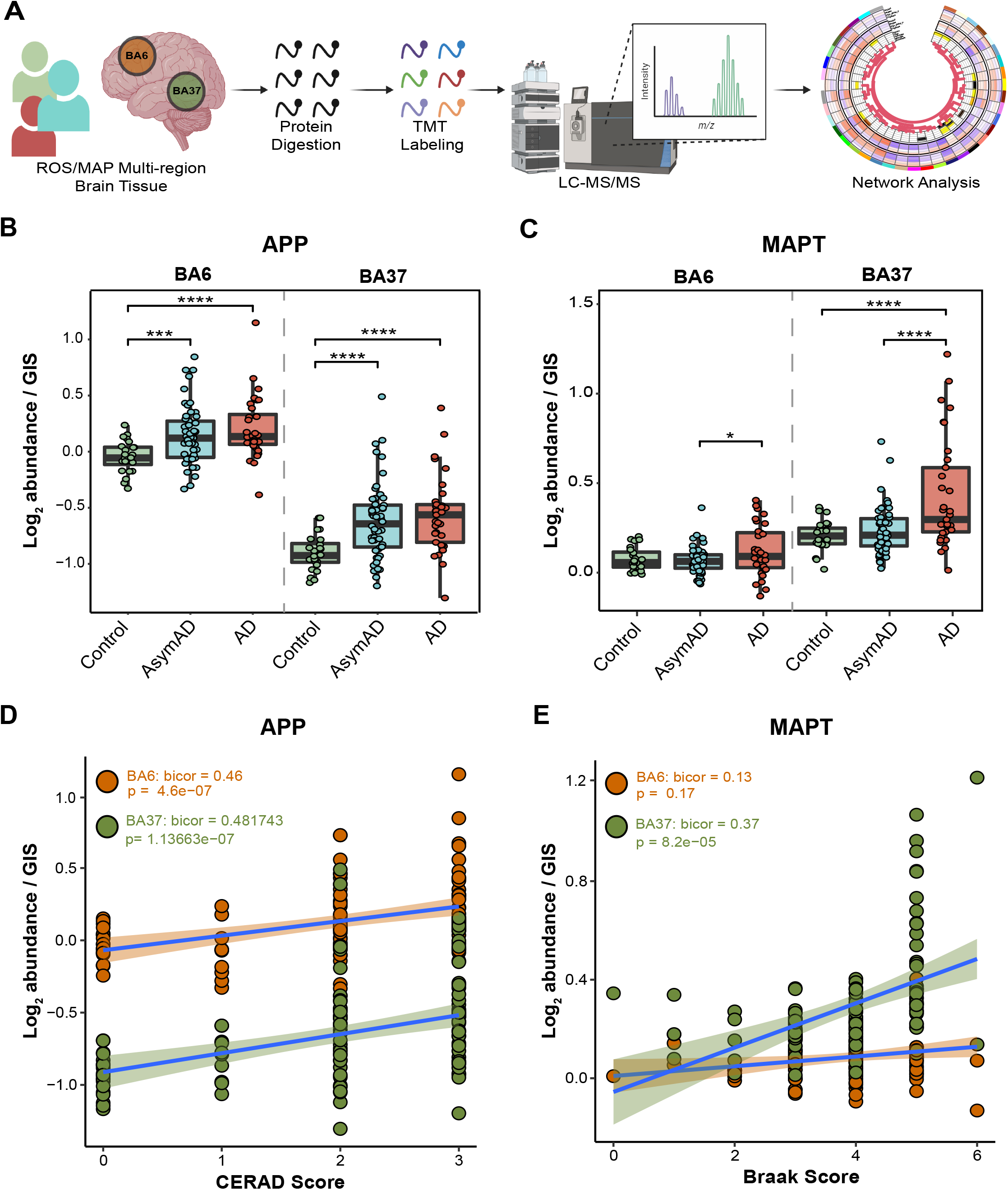
Proteomic measurements of amyloid and tau align with region-specific neuropathological burden. (**A**) Schematic representation of experimental workflow for matched human brain tissue samples across regions BA6 and BA37 from 109 ROSMAP cases that were enzymatically digested with trypsin into peptides and individually labeled with isobaric tandem mass tags (TMT) followed by LC-MS/MS. Log_2_ abundances were normalized as a ratio dividing by the central tendency of pooled standards (global internal standards, GIS) and median centered. Protein abundances were analyzed using differential and co-expression methods. (**B**) TMT-MS quantified APP normalized abundance is significantly increased in AsymAD and AD cases compared to Control. One-way ANOVA (BA6: F= 7.987, p=>0.001; BA37: F=9.469, p=>0.001) with Tukey’s multiple comparisons test. (**C**) TMT-MS quantified MAPT normalized abundance is significantly increased in AD. One-way ANOVA (BA6: F= 3.522, p=<0.05; BA37: F= 12.69, p=>0.001) with Tukey’s multiple comparisons test. (**D**) APP normalized abundance and CERAD scores positively correlate in each brain region. Biweight midcorrelation (Bicor) and pvalue (BA6: bicor=0.46, p=4.6e-07; BA37: bicor=0.482, p=1.1e-07). Best fit line for each region determined by linear model, confidence interval is shaded around line. (**E**) MAPT normalized abundance and Braak scores positively correlate in BA37. Bicor and pvalue (BA6: bicor=0.13, p=0.17; BA37: bicor=0.37, p=8.2e-05). Best fit line for each region determined by linear model, confidence interval is shaded around line. *p<0.05, ***p<0.005, ****p<0.001; F=F value; Bicor= biweight midcorrelation.

Prior to systems-level analyses, TMT-MS-quantified protein levels related to the primary AD pathologies, APP (amyloid precursor protein) and MAPT (microtubule associated protein tau), were compared across disease groups and brain regions (**Fig. 1B-C**). APP levels served as a surrogate measurement for Aβ and as expected were significantly higher in AsymAD and AD cases compared to Controls in both brain regions (**Fig. 1B**; (18, 21)). MAPT levels were significantly higher in AD compared to AsymAD in BA6 and significantly increased in AD compared to both Control and AsymAD in BA37 (**Fig. 1C**). APP levels positively correlated with CERAD scores, irrespective of brain region, and exhibited higher baseline levels in BA6 (**Fig. 1D**). MAPT levels positively correlated with Braak scores in BA37, but not BA6 (**Fig. 1E**). These measurements align with well-established region-specific neuropathological burden observed in AD in which amyloid pathology manifests initially in neocortical regions while tau tangle pathology originates and intensifies in trans-entorhinal cortex regions before spreading to temporal and frontal cortical areas (22, 23). Collectively, these strong positive correlations of APP levels with CERAD scores and MAPT levels with Braak scores highlight the accuracy of the proteomic measurements in quantifying relevant AD neuropathological burden.

Consistent with these targeted pathology-linked proteins (APP and MAPT), differential expression of all quantified proteins was assessed (**Supplemental Tables 1 and 2**). As expected, proportional numbers of significantly different proteins were identified in disease groups compared to controls (BA6: Control vs AsymAD = 222, Control vs AD = 1,102; BA37: Control vs AsymAD = 129, Control vs AD = 1,550), demonstrating differences in the proteome correspond proportionately with differences in neuropathology (21). In addition, the greater number of differentially expressed proteins in BA37 compared to BA6, where extent of pathology would be greater, further corroborates alignment of proteomic measurement and region-specific burden in AD.

### Regional brain co-expression network analysis reveals modules associated with AD pathology and cognition

A correlation network was constructed using the consensus weighted gene co-expression network analysis (cWGCNA) algorithm, a systems biology approach to identify biologically meaningful, co-expression patterns (24, 25). The consensus configuration allows the identification of highly preserved modules, or clusters of interconnected proteins, shared across BA6 and BA37 while retaining region-specific relationships (**Fig. S2B-C**). A total of 39 co-expression modules (M1-M39) were defined, ranging in size from 36 members (M39) as the smallest and 473 members (M1) as the largest (**Fig. 2A**; **Supplemental Tables 3 and 4**). Similar patterns of intermodule relationships were observed in BA6 and BA37 (**Fig. S2A**). Gene ontology (GO) analysis was performed on the protein members of each consensus module and top GO terms were considered representative of module biology. To detect modules related to neuropathological burden and cognitive changes, module eigenproteins (MEs) were correlated with Aβ plaque (“Amyloid”) and neurofibrillary tangle (NFT; “Tangles”) burden in the brain at autopsy as well as global cognitive scores and cognitive slope for each person prior to death (**Supplemental Tables 5 and 6**). Immunohistochemistry and systematic sampling of 8 brain regions were averaged to determine Amyloid and Tangle load (26). Global cognition is a composite score of 19 cognitive performance tests and cognitive slope is calculated based on changes in cognitive performance over time (12). To understand group-wise differences in MEs, modules were further characterized according to AD vs Control and AsymAD vs AD pairwise differences. Finally, cell-type contribution of each module was assessed by determining cell-type marker enrichment for neuronal, oligodendrocyte, astrocyte, microglia and endothelial cell-types (**Supplemental Tables 7 and 8**) (18).

**Figure 2.**
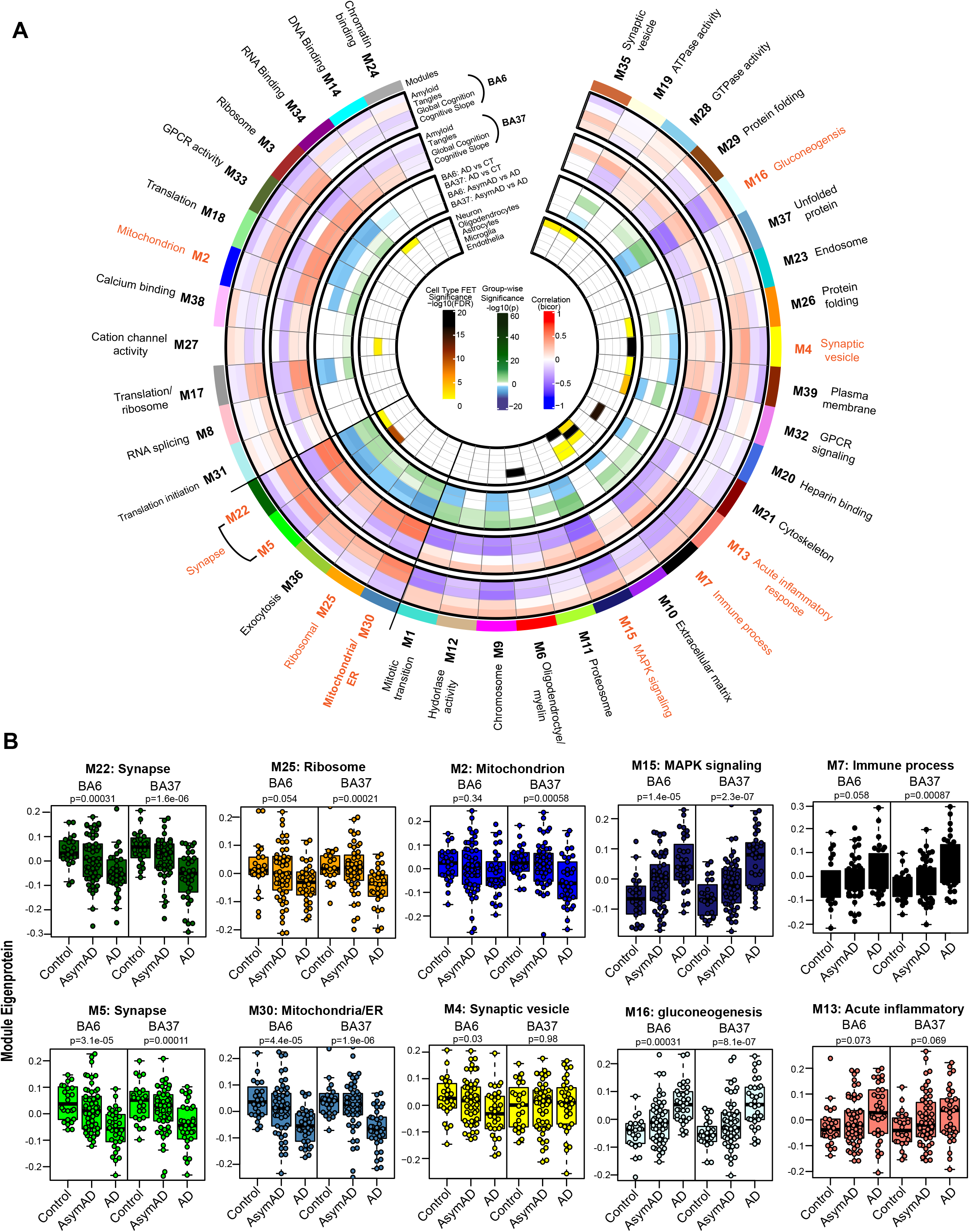
Consensus correlation network of a multi-region human brain proteome. (**A**) A consensus correlation network (cWGCNA) was constructed with 7,787 proteins across BA6 and BA37 and yielded 39 co-expression protein modules. In the inner-most heatmap, enrichment of cell-type markers (as determined by one-way Fisher’s exact test) for each module is visualized for neuronal, oligodendrocyte, astrocyte, microglial and endothelial cell-types. The panel outside of the cell-type results highlights group-wise differences in module eigenproteins for AD vs Control and AsymAD vs AD in each brain region. The two outer-most heatmaps depict the correlation (Bicor) of module eigenproteins with pathological (Amyloid and Tangle burden) and clinical (Global cognitive function and cognitive slope) phenotypes for both brain regions. Modules are identified by color and number, accompanied by top gene ontology (GO) terms representative of modular biology. Scale bars for cell-type enrichment (darker color indicates stronger enrichment), weighted group-wise eigenprotein difference (darker green correspond to stronger positive relationship and deeper blue indicates stronger negative relationship) and bi-directional module— trait relationships (red indicating positive correlation and blue indicating negative correlation) are at the center of the plot. (**B**) Module eigenproteins (MEs) grouped by diagnosis (Control, AsymAD and AD) were plotted as box and whisker plots for modules of interest, chosen based on their preservation in AsymAD compared to AD and relationships to cognitive measures. MEs were compared in each brain region using one-way ANOVA, unadjusted pvalues are shown. Box plots represent median, 25th and 75th percentiles. Box hinges represent the interquartile range of the two middle quartiles with a group. Error bars are based on data points 1.5 times the interquartile range from the box hinge.

Module preservation of the current consensus regional network was compared to a recent, large-scale network analysis of human dorsolateral prefrontal cortex (Brodmann area 9, BA9) generated from ROSMAP and Banner cases (18). Notably, approximately 95% (37/39) of the consensus modules preserved with the previous TMT-MS network (**Fig. S3A**) and all 39 consensus modules from this study had significant protein overlap with at least one module from the BA9 network (**Fig. S3B**). Thus, modules generated by cWGCNA are robust and highly preserved across different cohorts and brain regions.

From the present network, we observed interplay of module biology with cognition, disease status and brain region. Specifically, a cluster of five modules were identified as positively correlated with cognition and were increased in AsymAD compared to AD similarly in both brain regions: M22 Synapse, M5 Synapse, M36 Exocytosis, M25 Ribosome, M30 Mitochondria/ER (**Fig. 1**). We also observed modules following this pattern in only one brain region: M2 Mitochondrion was only significant in BA37 and M4 Synaptic vesicle was only significant in BA6. In contrast, M15 MAPK signaling and M16 Gluconeogenesis were the most strongly negatively correlated with cognition and decreased in AsymAD compared with AD. These findings are consistent with previous proteomic findings in BA9 where sugar metabolism and MAPK signaling modules were significantly related to cognition (18). Overall, we generated a consensus network, highly preserved with previous brain proteome network modules, that sufficiently outlined key differences and an interrelationship between clinical traits, disease groups and even regional brain differences.

### Nomination of resilience-associated modules and NRN1 as top protein candidate

To increase external validity and nominate modules linked to resilience in an unbiased manner, we integrated results from a recent brain proteome-wide association study (PWAS) of cognition that evaluated the association of cortical protein abundances with cognitive resilience from an independent TMT-MS proteomic analysis of ROSMAP tissues adjusted for AD pathologies (27). Higher abundances of proteins related to slower rates of cognitive decline were considered to confer greater resilience while higher abundance of proteins associated with faster rate of cognitive decline were considered to confer less resilience. Four modules were identified as significantly enriched with proteins conferring greater cognitive resilience: M22 Synapse, M5 Synapse, M36 Exocytosis and M30 Mitochondria/ER (**Fig. 3A; Supplemental Table 9**). In addition, four modules were found to be significantly enriched for proteins conferring less cognitive resilience: M11 Proteosome, M15 MAPK signaling, M32 GPCR signaling and M16 Gluconeogenesis (**Supplemental Table 10**). Of the modules associated with greater resilience, the protein constituents of M5 and M22 were strongly representative of synaptic biology and enriched for neuronal markers. To further confirm association of M5 and M22 to cognitive preservation, MEs were correlated to cognitive slope and indicated strong positive correlation in both brain regions (**Fig. 3B**). Consistently, differential expression comparing AsymAD with AD of proteins specific to M5 and M22 exhibited a strong bias towards an increase or upregulation of proteins in AsymAD (**Fig. 3C**). Two proteins significantly upregulated in AsymAD were also significant in the aforementioned PWAS of cognition, Neuritin (NRN1) and Rabphilin-3A (RPH3A). NRN1 was more significantly differentially expressed than RPH3A and the most significant protein associated with increased cognitive resilience in the PWAS. NRN1 abundance in both brain regions was compared across disease groups and indicated preserved levels in AsymAD similar to controls but was significantly downregulated in AD (**Fig. 3D**). NRN1 abundance also strongly, positively correlated with cognitive measures including global cognition and cognitive slope (**Fig. 3E-F**). Furthermore, variance partition analysis of global cognition in BA6 and BA37 identified NRN1 as the top (B6: ~26% variance explained) and second (BA37: ~38% variance explained) protein explaining the highest variance in global cognition (**Fig. S4A-B; Supplemental Tables 11 and 12**).

**Figure 3.**
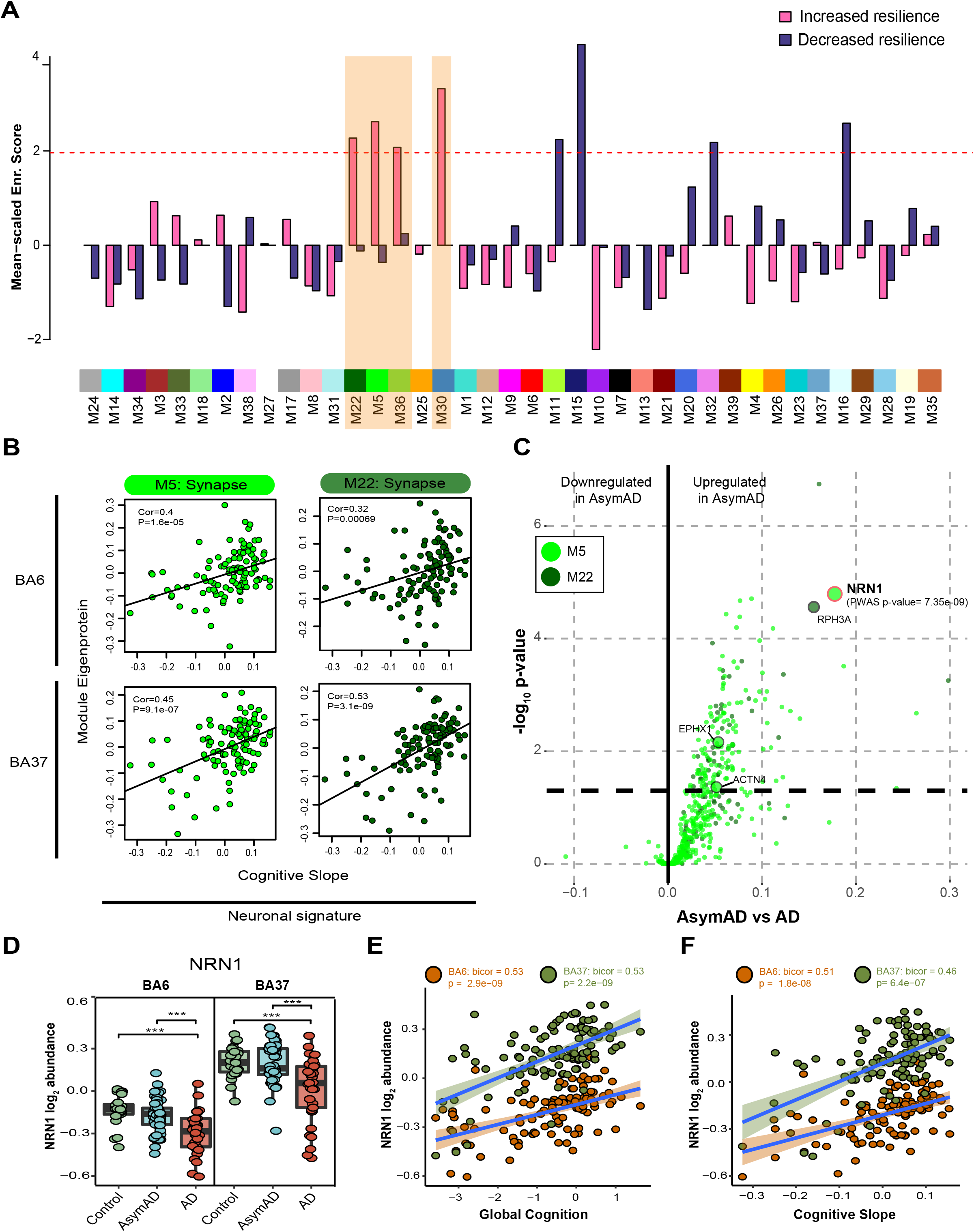
Integrated proteomics of human brain reveals NRN1 as a top resilience candidate. (**A**) Significant enrichment of modules associated with increased cognitive resilience were identified by PWAS in consensus modules. The dashed red line illustrates the significance cutoff corresponding to a Z score of 1.96 or p=0.05. Significant, increased resilience modules are highlighted in orange. (**B**) PWAS significant, synaptic modules M5 and M22 positively correlate with cognitive slope, irrespective of brain region. Bicor and pvalues (BA6: M22 cor=0.32, M22 p=0.00069, M5 cor=0.4, M5 p=1.6e-05; BA37: M22 cor=0.53, M22 p=3.1e-09, M5 cor=0.45, M5 p=9.1e-07). (**C**) Differential expression comparing AsymAD and AD groups from M5 and M22 module members. Protein fold-change is the x-coordinate and the -log_10_ pvalue from one-way ANOVA is the y coordinate for each protein. Proteins above the dashed line (p=0.05) are considered significantly differentially expressed. Large circles highlight proteins that were significant by PWAS (α=5e-06). (**D**) NRN1 abundance is significantly reduced in AD. One-way ANOVA (BA6: F=13.25, p=<0.001; BA37: F=13.68, p=<0.001) with Tukey test. (**E**) NRN1 abundance correlates positively with global cognitive performance. Bicor and pvalues (BA6: bicor=0.53, p=2.9e-09; BA37: bicor=0.53, p=2.2e-09). (**F**) NRN1 abundance correlates positively with cognitive slope. Bicor and pvalues (BA6: bicor=0.51, p=1.8e-08; BA37: bicor=0.46, p=6.4e-07).

In summary, integration of independent human proteomic data identifies protein modules associated with cognitive resilience across two brain regions. Four modules significantly enriched for proteins suggested to promote greater cognitive resilience in life, two of which captured biology found to be vulnerable in AD yet preserved in asymptomatic cases – synaptic integrity. Among the hub proteins of these modules, NRN1 levels were highly up-regulated and preserved in AsymAD compared to AD. NRN1 co-expression with synaptic biology-linked proteins and positive correlation with cognition further supports the hypothesis that NRN1 is a molecular effector of cognitive resilience.

### NRN1 prevents Aβ_42_-induced dendritic spine degeneration

The preservation of dendritic spines is hypothesized to maintain memory and information processing in resilient patients who harbor high levels of Aβ pathology but are cognitively normal (9, 28). Numerous studies indicate that Aβ can induce dendritic spine degeneration in cellular and animal models of AD (29–32). Henceforth, protecting spines from Aβ represents a rational therapeutic strategy to promote resilience and delay dementia onset. Past studies provided evidence that NRN1 exists predominantly as a soluble form *in vivo* and exerts neurotrophic effects on synaptic maintenance and neuronal survival (33–35). To test whether NRN1 is protective against Aβ_42_-induced dendritic spine degeneration, rat hippocampal neurons were isolated at E18 and cultured at high-density on glass coverslips. To visualize dendritic architecture, neurons were transiently transfected with a plasmid encoding Lifeact-GFP at DIV 14. Cultures were treated with NRN1 or co-treated with NRN1 and Aβ_42_ oligomers for 6 hours, then fixed, and processed for widefield microscopy and subsequent neuronal three-dimensional reconstructions for dendritic spine morphometric analysis (**Fig. 4A**). Consistent with previous reports (36), spine density was reduced significantly after exposure to Aβ_42_ in comparison to DMSO controls, however co-treatment with NRN1 prevented Aβ_42_-induced spine degeneration (**Fig. 4, B and C**). Examination of dendritic spine morphologic subclasses revealed that Aβ_42_ exposure significantly decreased thin spine density in comparison to DMSO controls, but these detrimental effects were blocked in the presence of NRN1 (**Fig. 4D**). Notably, the proportion of thin spines were increased with NRN1 treatment compared to DMSO, while Aβ_42_ promoted an increase in the proportion of dendritic filopodia (**Fig. 4E**). Exposure to Aβ_42_ and/or NRN1 did not significantly alter dendritic spine length or head diameter in comparison to DMSO controls (**Fig. 4, F and G** and **Fig. S5, A and B**). These findings suggest that NRN1 can protect against Aβ_42_-induced dendritic spine loss.

**Figure 4.**
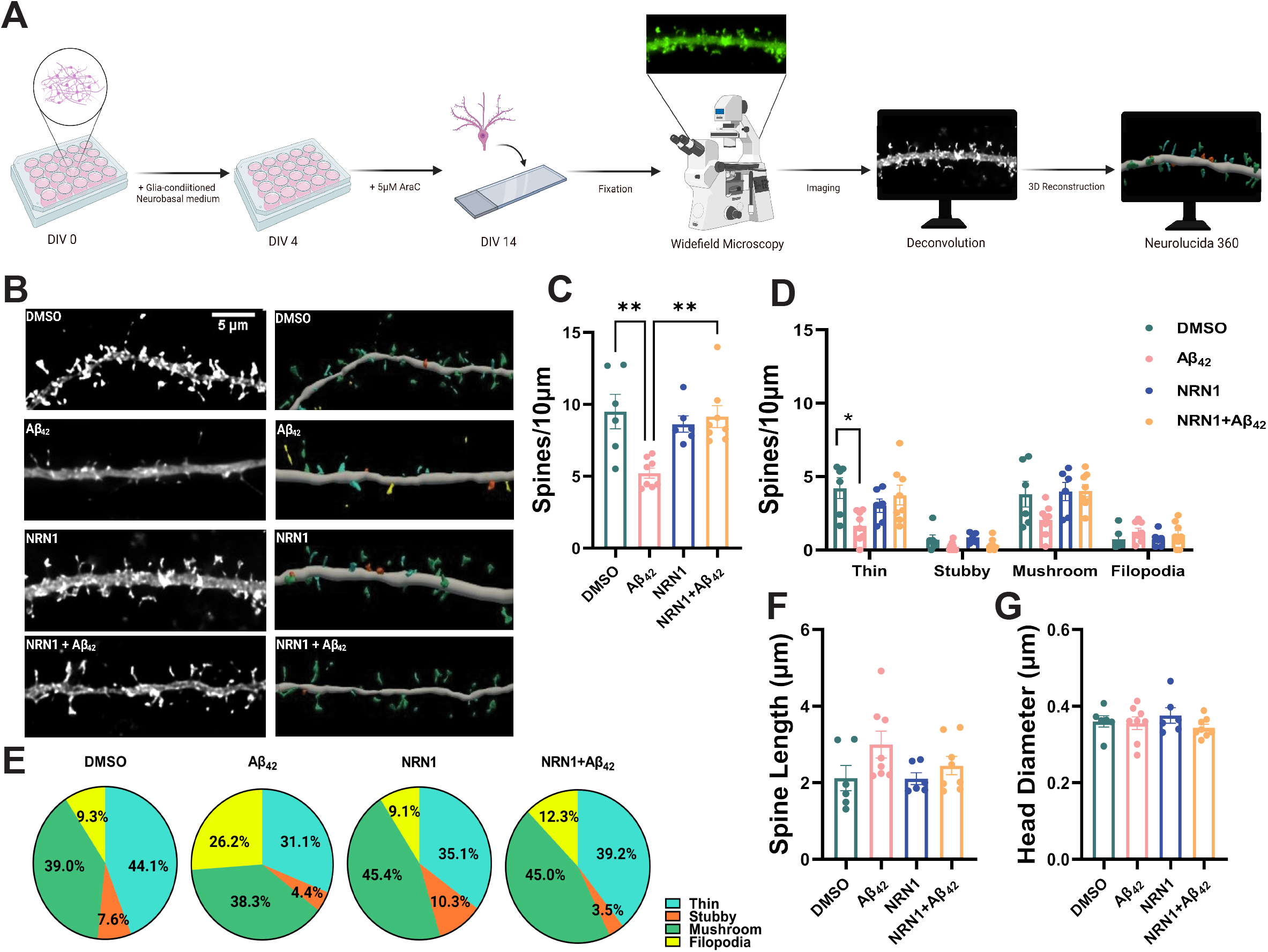
Aβ_42_-induced dendritic spine degeneration is blocked by NRN1. (**A**) Schematic representation of primary rat hippocampal neuron treatment and dendritic spine morphometric analysis. (**B**) Representative maximum-intensity wide-field fluorescent images of hippocampal neurons after deconvolution (left). Corresponding three-dimensional reconstructions of dendrites generated in Neurolucida 360 (right), with dendritic spines color-coded by spine type (blue = thin, orange = stubby, green = mushroom, yellow = filopodia). Scale bar, 5 μm. N = 6 to 8 neurons (one dendrite per neuron) were analyzed per experimental condition. (**C**) Dendritic spine type frequency in hippocampal neurons exposed to DMSO, 500nM Aβ_42_, 150 ng/mL NRN1, or 150 ng/mL NRN1 and 500nM Aβ_42_. (**D**) Dendritic spine density in hippocampal neurons exposed to DMSO, 500nM Aβ_42_, 150 ng/mL NRN1, or 150 ng/mL NRN1 and 500nM Aβ_42_. Data are means + SEM. **P < 0.01 (DMSO vs. Aβ_42_, actual P = 0.0025) (Aβ_42_ vs NRN1+Aβ_42_, actual P = 0.0026) by one-way ANOVA with Tukey’s test. *P < 0.05 (Aβ_42_ vs NRN1, actual P = 0.0177) by one-way ANOVA with Tukey’s test. (**E**) Overall mean of dendritic spine length and (**F**) head diameter. Data represent the mean + SEM. Related data are shown in **Fig. S1**. (**G**) Dendritic spine density of thin, stubby, or mushroom spines per 10 μm. Data are means + SEM. *P < 0.05 (DMSO vs. Aβ_42_, actual P = 0.0218) by one-way ANOVA with Tukey’s test. (Thin, Aβ_42_ vs NRN1+Aβ_42_, actual P = 0.0501) (Mushroom, DMSO versus Aβ_42_, actual P = 0.1514) (Mushroom, Aβ_42_ versus NRN1+Aβ_42_, actual P = 0.0598) by one-way ANOVA with Tukey’s test.

To exclude the possibility that NRN1 directly binds to soluble Aβ_42_ oligomers and in turn neutralizes each protein’s independent effects in primary neurons, we performed an *in vitro* amyloid aggregation assay. Human recombinant Aβ_42_ fibrilization was measured by thioflavin T (ThT) fluorescence in the presence or absence of NRN1 for 20 consecutive hours (**Fig. S6A**). There were no distinguishable differences in self-assembly and aggregation between Aβ_42_ alone and Aβ_42_ and NRN1 together. Following the fluorometric assay, soluble and pellet fractions of assay products were probed via western blot and silver stain (**Fig. S6B-C**). Nearly all NRN1 immunoreactivity was detected in the soluble fraction whereas Aβ was primarily concentrated in the pelleted fraction. Importantly, the molar concentration of Aβ_42_ was 10-fold greater in the aggregation assay and the molar concentration of NRN1 was >30-fold greater, suggesting that even at very high concentrations NRN1 does not impede Aβ_42_ fibrilization. These studies suggest that NRN1’s protection of spines in the presence of Aβ_42_ was unlikely due to an artifact of NRN1 and Aβ_42_ directly binding *in vitro* to prevent individual proteins from interacting with dendritic spines.

### NRN1 protects against Aβ_42_-induced neuronal hyperexcitability

Aβ-induced dendritic degeneration and spine loss cause reductions in the overall area and volume of neurons, rendering them more electrically compact(29, 32). These detrimental effects induce neuronal hyperexcitability, which consequently drives abnormal circuit synchronization and cognitive impairment in AD mouse models and patients(29, 37, 38). To test whether NRN1 is protective against Aβ_42_-induced neuronal hyperexcitability, we seeded rat primary hippocampal neurons on microelectrode arrays (MEAs) and performed baseline recordings at DIV 14. Action potential frequency, referred to as mean firing rate, was measured to assess neuronal excitability. Immediately after the baseline recording, neurons were exposed to DMSO, Aβ_42_, NRN1, and/or NRN1 plus Aβ_42_ for 6 hours followed by a second recording (**Fig. 5A**). DMSO did not increase mean firing rates in comparison to baseline (**Fig. 5, B and C**). Consistent with previous findings(32), Aβ_42_ significantly increased mean firing rates in comparison to baseline (**Fig. 5, B and D**). While NRN1 significantly increased mean firing rates in comparison to baseline, simultaneous exposure to Aβ_42_ and NRN1 was comparable to baseline (**Fig. 5, B, E, and F**). The total number of active neurons per experimental group could not account for the effects on mean firing rates (**Fig. S7**). Collectively, these results indicate that the dendritic spine resilience provided by NRN1 is protective against Aβ_42_-induced hyperexcitability.

**Figure 5.**
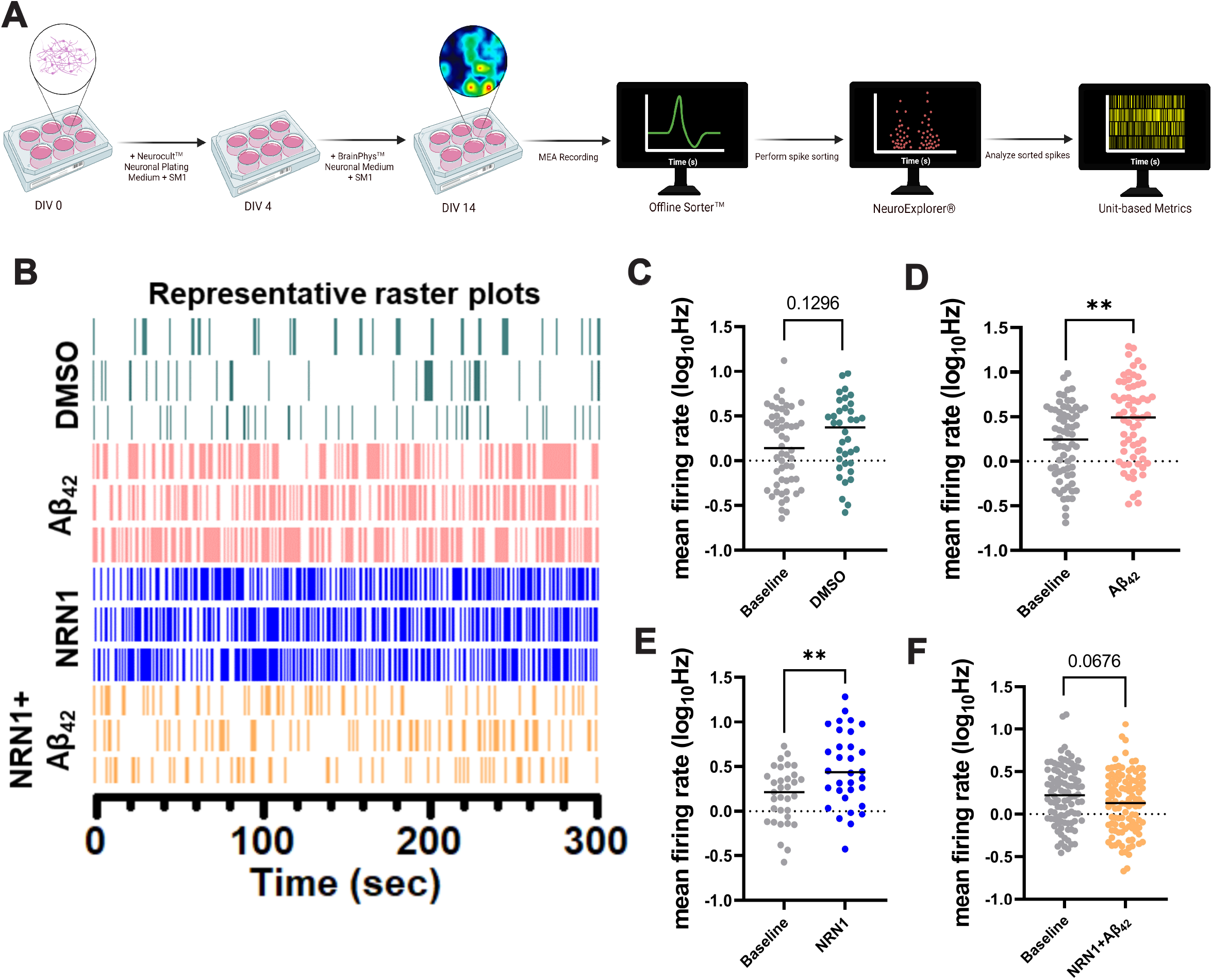
NRN1 protects against Aβ_42_-induced neuronal hyperexcitability. (**A**) Schematic representation of primary rat hippocampal neuron treatment and single neuron electrophysiology analysis. (**B**) Representative raster plots from three units after exposure to DMSO, 500nM Aβ_42_, 150 ng/mL NRN1, or 150 ng/mL NRN1 and 500nM Aβ_42_. (**C**) Mean firing rate at DIV14 in hippocampal neurons treated with DMSO, compared to baseline (n = 36-54 neurons, unpaired Student’s t test; p = 0.1296). (**D**) Mean firing rate at DIV14 in hippocampal neurons treated with 500nM Aβ_42_, compared to baseline (n = 65-68 neurons, unpaired Student’s t test; p = 0.0022). (**E**) Mean firing rate at DIV14 in hippocampal neurons treated with 150 ng/mL NRN1, compared to baseline (n = 32-33 neurons, unpaired Student’s t test; p = 0.0023). (**F**) Mean firing rate at DIV14 in hippocampal neurons treated with 150 ng/mL NRN1 and 500nM Aβ_42_, compared to baseline (n = 100-107 neurons, unpaired Student’s t test; p = 0.0676).

### NRN1 treatment alters the neuronal proteome in cultured neurons

Although NRN1 has been well characterized for roles in synaptic plasticity and maturation, the receptor(s) and downstream signaling events enabling neuronal functions of NRN1 remain poorly understood. To identify proteins and broader pathways impacted by NRN1 treatment, rat primary cortical neurons were treated with NRN1 recombinant protein at DIV14 for 6 hours at the same concentration as previously tested in dendritic spine and MEA assays (**Fig. 6A**). Following treatment, cells were lysed and prepared for TMT-MS analysis. A total of 8,238 proteins were quantified and used for differential expression analysis (**Supplemental Table 13**). Comparing NRN1 treated and vehicle treated neurons, 445 proteins were significantly increased and 400 proteins were significantly decreased following NRN1 treatment (**Fig. 6B**; **Supplemental Table 14**). As expected, NRN1 was identified among proteins significantly increased in the NRN1 treatment group. GO analysis of significantly changed proteins found strong bias of synaptic and cell projection functions upregulated with NRN1 exposure (**Fig. 6C**). In addition, proteins involved in functions related to oxidation and metabolic processes were decreased following NRN1 treatment. These results support previously observed functions of NRN1 in promoting synaptic function (15), provide a reference of downstream and coregulated proteins impacted by incubation with exogenous NRN1 and allow the inference of molecular mediators driving neuronal firing and synaptic density changes observed in our dendritic spine morphometric and MEA analyses. Furthermore, pathways decreased following NRN1 treatment were related to metabolism and cellular energetics, systems often dysregulated and increased in AD, implicating NRN1 as a dual-action molecular effector able to increase proteins typically vulnerable to or lost in AD and to decrease proteins aberrantly increased in disease.

**Figure 6.**
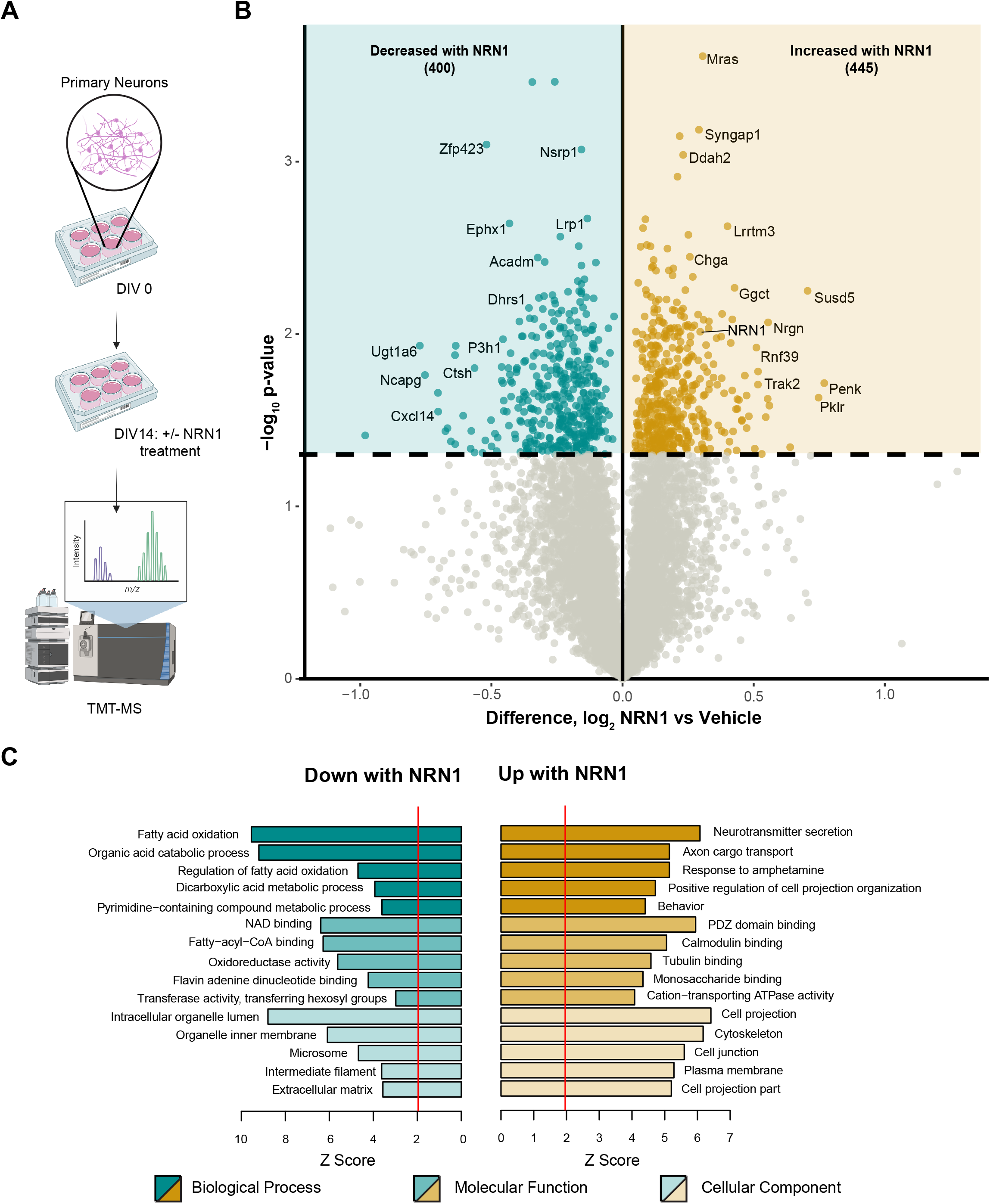
NRN1 treatment induces changes in the neuronal proteome related to broad synaptic functions. (**A**) Schematic representation of rat primary cortical neuronal culture workflow in which neurons were maintained in neurobasal medium for 14 days, treated with 150ng/mL of NRN1 and analyzed via TMT-MS. (**B**) Differential protein expression between NRN1 treated and vehicle treated neurons (n=8,238 proteins). Proteins above the dashed line (p=0.05) are considered significantly differentially expressed. Student’s t-test was used to calculate pvalues. (**C**) Gene ontology of significantly differentially expressed proteins in NRN1 treated neurons. A Z-score above 1.96 was considered significant (p<0.05).

### NRN1 engages protein targets linked to cognitive resilience in human brain

The ability of a nominated resilience protein candidate to engage relevant human biology is critically important to the translational value of the target. Therefore, we applied an integrative analysis to resolve where NRN1-driven changes observed in the rat neuronal proteome mapped to the human brain proteome. Significance of protein overlap between the rat neuronal proteome and individual modules within the human brain consensus network was determined by one-tailed Fisher’s exact test (**Fig. 7A; Supplemental Table 15**). The rat neuronal proteomic data was subset into three for this analysis and included: I) all proteins quantified, II) only those that were significantly upregulated by NRN1 treatment and III) only those that were significantly downregulated by NRN1 treatment. This analysis revealed 17 of the 39 modules with statistically significant overlap from rat neuronal proteins into the human brain network (**Fig. 7A** -top row, p<0.05). An additional 7 modules overlapped with rat neuronal proteins significantly differentially expressed with NRN1 treatment (middle and bottom rows). M22 Synapse, M5 Synapse, M4 Synaptic vesicle and M19 ATPase activity in the human brain network were enriched for proteins increased following NRN1 treatment. Human modules M8 RNA splicing, M31 Translation initiation and M12 Hydrolase activity were enriched for proteins decreased following NRN1 treatment. The majority of proteins significantly upregulated by NRN1 treatment in the rat neuronal proteome overlapped with human modules M5 and M22 (**Fig. 7B**). Notably, M5 and M22 are enriched with neuronal markers and identified as top resilience-associated modules (**Fig. 3A-B**). Further, nearly all proteins increased by NRN1 in the rat neuronal proteome were significantly increased in the human asymptomatic cases (**Supplemental Table 16**) and significantly correlated with cognitive slope (58 out of 83 or ~70%, pvalue≤0.05; **Supplemental Table 17**). Importantly, VGF (VGF nerve growth factor inducible) was present in this overlap and has been nominated as a potential therapeutic target of greatest interest by AD research working groups; predictably its changes in brain and cerebrospinal fluid (CSF), which correlate strongly with cognition, have been described as a causal driver of AD pathophysiology (39–41). These findings indicate that NRN1 target engagement is highly relevant to human resilience mechanisms identified by unbiased systems-level analyses and further support NRN1 as a bimodal mediator of physiological resilience.

**Figure 7.**
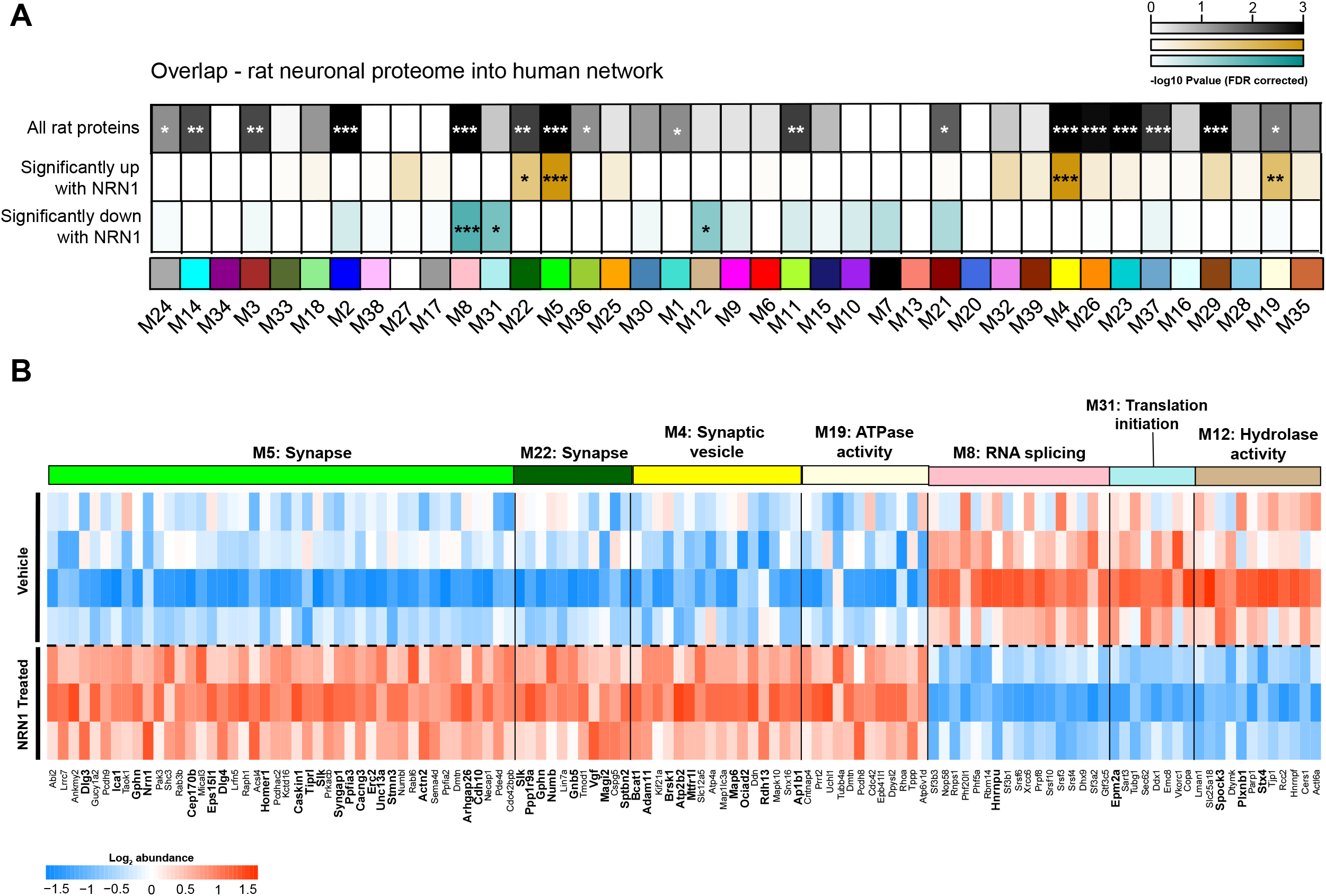
NRN1 engages proteins within modules linked to cognitive resilience in human brain. (**A**) To directly compare NRN1-induced changes in the context of human biology, a Fisher’s exact test was used to calculate significant enrichment of proteins from the entire rat proteome (top row), significantly upregulated with NRN1 (middle row) and significantly downregulated with NRN1 (bottom row) treatment across the 39 human consensus modules (0.05 > p > 0.01 = *, 0.01 > p > 0.005 =**, p < 0.005 = ***). (**B**) Proteins significantly impacted by NRN1 treatment that overlap with human modules were visualized as a heatmap. Rat protein abundance was compared using Bicor across NRN1 treated and vehicle treated groups.

## Discussion

In the present study, we implemented an integrative pipeline that pairs systems-level nomination of resilience-associated proteins in human brain with experimental mechanistic validation in a primary cell model. This approach enables both unbiased identification and bidirectional integration of molecular and clinical data. Following the current framework, TMT-MS based proteomic data from two independent studies and a total of three brain regions were incorporated to nominate communities of proteins strongly related to cognitive resilience in an unbiased manner. Modules associated with cognitive resilience were mined to identify a hub protein, NRN1, that was functionally validated for synaptic resilience against Aβ. To define overlapping neurobiology between NRN1’s effects on primary neurons and humans, TMT-MS proteomic data from the model system was fed back into the human brain proteome to identify convergent pathways relevant for resilience.

Correlation networks have been applied successfully to many biological and translational questions and demonstrated validity in identifying candidate biomarkers and therapeutic targets, including in ROSMAP (24, 42). Herein, cWGCNA resolved 39 co-expression modules across two brain regions from Control, AsymAD and AD cases. Applying the consensus configuration of WGCNA for matched brain tissues from the same cases identified protein communities shared across both BA6 and BA37. Module eigenprotein correlation with pathological and clinical traits further illuminated patterns of preservation in asymptomatic cases related to synaptic biology, cellular energetics and protein translation. Importantly, the majority of modules identified in this study preserve with a recent, large-scale network analysis of over 1000 cases from multiple institutions, supporting the strength and reproducibility of our findings (18). Results from an independent proteome-wide association study of cognition were then integrated for unbiased nomination of resilience associated modules, which included four modules significantly enriched for proteins conferring increased resilience. Among these, M5 and M22 were the most significantly enriched for synaptic biology and displayed strong positive correlation with cognitive performance in life. NRN1, a hub of M5, has been identified as a top protein candidate of resilience, previously by its relationship with cognitive trajectory (27) and in the current study by its preservation in AsymAD cases and correlation with elevated cognitive function in life. NRN1, also known as candidate plasticity gene 15 (*CPG15*), is a neurotrophic factor that was initially discovered in a screen to identify genes involved in activity-dependent synaptic plasticity in the rat dentate gyrus (43). Over the past two decades, the role of NRN1 in regulating neurodevelopment, specifically the formation of axonal arbors and dendritic branching, has been extensively studied (13, 14, 33, 44–48). In adult brain, NRN1 strongly correlates with synaptic maturation, long-term stability and activity-related plasticity (13, 14, 49) (15) (13, 48, 50). Importantly, NRN1 was identified among proteins previously shown in multiple studies to relate to increased cognitive function and resilience to AD, including VGF, NPTX2, and RPH3A (18, 51, 52). The established link between synaptic loss and cognitive impairment in AD, and given the predominance of synaptic proteins in our top resilience-associated modules, examining the impact of NRN1 on synaptic integrity and maintenance is foundational to determining its role in resilience.

Dendritic spines are small actin-rich protrusions off dendrites that serve as the postsynaptic sites of the majority of excitatory synapses in the brain. Spines exhibit remarkable variability in size, shape, and density along the length of dendritic branches (53–55). Spine structure is inseparably linked to spine function and spines are classified on the basis of their three-dimensional morphology as stubby, mushroom, thin or filopodia (56, 57). Cognitive decline associated with aging is hypothesized to be driven by subtle alterations in dendritic spine density and morphology in mammals. Thin spine loss occurs with age in the dorsolateral prefrontal cortex and correlates with worsening cognitive performance (28, 58–60). In parallel, patients with AD can exhibit high rates of epileptic seizure activity which is associated with accelerated cognitive decline (37, 38, 61). In APP transgenic mice, epileptiform activity is an indicator of network hyperexcitability which is driven by degeneration of hippocampal pyramidal neurons’ dendrites and dendritic spines (29). Loss of dendrites and spines reduces the total surface area of the cell and renders the neuron more electrically compact. In a compact neuron, synapse currents are translated more frequently which leads to increased action potential output, consequently inducing neuronal hyperexcitability and aberrant circuit synchronization (62). Similar to APP transgenic mice, exogenously applied Aβ_42_ oligomers can induce dendritic spine degeneration which subsequently causes hyperexcitability in cultured rodent hippocampal neurons (36). Using highly optimized three-dimensional modeling of spines in combination with MEA analyses, we show that exogenously delivered NRN1 protects against Aβ_42_-induced spine degeneration and hyperexcitability. Moreover, our results indicate that in cultures treated with NRN1 alone, alterations in spine density or morphology were not observed. However, NRN1 alone increased mean action potential firing rates. The mechanisms by which NRN1 increases action potential frequency are not due to alterations in spine density or structure, unlike the effects of Aβ_42_. We posit that the elevation in mean firing rates are due to NRN1-mediated modification of the synaptic proteome, which are highlighted by increases in protein abundance from M4, M5, and M22 from human brain (**Fig. 7**). Yet, it remains to be determined whether the downstream pathways of NRN1-protection against Aβ_42_ are similar to or different from how NRN1 affects the proteome in the absence of Aβ_42_. Notably, Choi et al. showed that over-expression of NRN1 in cultured hippocampal neurons increased mini excitatory postsynaptic current frequency, which mirrors our findings that NRN1 alone increased action potential firing rate (63). Furthermore, electrophysiology studies by An et al. demonstrated that brain infusion of recombinant NRN1 (similar to the reagents used in this study) into Tg2576 APP transgenic mice rescued deficits in hippocampal long-term potentiation in the Schaffer collateral pathway (48). Collectively, these findings support the promise of NRN1 as a therapeutic target to support synaptic mechanisms of resiliency in preclinical stages of AD.

The ROSMAP studies are information-rich longitudinal aging studies that have invaluably contributed to understanding the complexity of aging and disease-related changes over time. However, this cohort is primarily made up of non-Latino white participants and historically lacks equal representation from diverse populations. Recent reports indicate Black and Hispanic populations are disproportionately more likely to have AD compared to older white Americans (64), which highlights a potential limitation of the current study. In addition to population demographics, the use of multiple definitions of resilience and how researchers identify this group adds complexity to generalizable interpretation of findings (7, 8). The current study used the combination of pathological and cognitive metrics to differentiate asymptomatic from symptomatic cases by imposing cutoffs which would identify resilient cases with the greatest confidence, however, there may be more to learn from cases not captured by this strategy. Another potential limitation of the current study is that NRN1 neuroprotection was only assessed for Aβ insult and not tau. Quantitative neuropathological studies indicate that asymptomatic cases typically have lower levels of tau pathology but comparable levels of amyloid burden in the brain at autopsy compared to symptomatic AD cases (10). Thus, understanding the impact of NRN1 on Aβ insult is highly relevant to the pathological context observed in resilient brains. Future work investigating the interaction or effects of NRN1 on tau neuropathology may provide additional insights into NRN1 neuroprotection relevant to at-risk populations.

Conventional benchtop-to-bedside strategies for identifying therapeutic targets have generated an abundance of data in clinical trial settings, but unfortunately often fail. Reverse translation, or bedside-to-benchtop, begins with human observational studies and works backwards to pinpoint potential mechanisms and therapeutic targets for investigation. This paradigm allows information from clinical and laboratory settings to follow a cyclical process instead of a linear one, and thereby is tunable and more likely to lead to successful clinical interventions (65). In the current study, we use human postmortem brain proteomic data with incorporated antemortem clinical phenotypic data (e.g., cognitive trajectory in life) to unbiasedly nominate protein modules important for resilience. NRN1 was targeted in this analysis and validated for neuroprotective efficacy in a neuronal model system. Finally, findings from our experimental models were re-integrated back into our human data to generate a distinct collection of proteins and associated biology linked to cognitive resilience in humans with high confidence. Overall, this study established an integrative, non-linear pipeline for the identification and validation of resilience-associated proteins similarly to the reverse translation paradigm. The current work provides a valuable framework for investigating molecular and physiological underpinnings of resilience directed from patient samples and cognitive changes in life.

## Data and Code Availability

Raw mass spectrometry data from the frontal and temporal cortex, study background and additional metadata on the ROSMAP cognitive resilience study can be found at https://www.synapse.org/#!Synapse:syn22695346. Pre- and post-processed protein expression data and case traits related to this manuscript are available at https://www.synapse.org/#!Synapse:syn25006620. The results published here are in whole or in part based on data obtained from the AMP-AD Knowledge Portal (https://adknowledgeportal.synapse.org). The AMP-AD Knowledge Portal is a platform for accessing data, analyses and tools generated by the AMP-AD Target Discovery Program and other programs supported by the National Institute on Aging to enable open-science practices and accelerate translational learning. The data, analyses and tools are shared early in the research cycle without a publication embargo on secondary use. Data are available for general research use according to the following requirements for data access and data attribution (https://adknowledgeportal.synapse.org/#/DataAccess/Instructions). Additional ROSMAP resources can be requested at www.radc.rush.edu. Raw mass spectrometry and processed abundance data for the rat neuronal proteome is available at https://www.synapse.org/ADresilienceRat.

## Methods and Materials

### Chemicals and Reagents

For primary neuron experiments, Aβ_42_ oligomers were purchased from Bachem and prepared as previously described (66). Aβ_42_ was resuspended in 1X Hanks’ balanced salt solution (HBSS) and Dimethyl sulfoxide (DMSO) then placed in 4°C overnight. Recombinant human Neuritin protein (Abcam, ab69755) was reconstituted in water to a concentration of 0.1 mg/mL. For the Thioflavin T (ThT) aggregation assay, recombinant human Aβ_42_ (5 μM) (rPeptide, # A-1170-1) was handled essentially as described (67) and detailed below. Plasmid encoding Lifeact-GFP was a generous gift from Dr. Gary Bassell, Emory University School of Medicine, Atlanta, GA, USA.

### Human postmortem brain tissue and case classification

Paired brain tissue samples from frontal cortex (Brodmann area 6, BA6) and temporal cortex (Brodmann area 37, BA37) were obtained from the Religious Orders Study and Rush Memory and Aging Project (ROS/MAP; n=256 total samples) in accordance with proper Institutional Review Board (IRB) protocols. Postmortem neuropathological evaluation of neuritic plaque distribution was performed according to the Consortium to Establish a Registry for Alzheimer’s Disease (CERAD) criteria (68) and extent of neurofibrillary tangle pathology was assessed with the Braak staging system (23). All case metadata are available at https://www.synapse.org/#!Synapse:syn22695346. Case classification was determined according to a previously established and peer-reviewed strategy (18, 19). In brief, cases with CERAD scores of 0-1 and Braak scores 0-3 without dementia at last evaluation were defined as control (if Braak equals 3, then CERAD must equal 0); cases with CERAD scores 1-3 and Braak scores 3-6 without dementia at last evaluation were defined as AsymAD; cases with CERAD 2-3 and Braak 3-6 with dementia at last evaluation were defined as AD. Dementia was defined at MMSE (Mini Mental State Examination) scores <24 (69).

### Brain tissue homogenization

Sample homogenization was performed as previously descried (18). Approximately 100 mg (wet tissue weight) of brain tissue was homogenized in 8 M urea lysis buffer (8 M urea, 10 mM Tris, 100 mM NaH_2_PO_4_, pH 8.5) with HALT protease and phosphatase inhibitor cocktail (Thermo Fisher Scientific) using a Bullet Blender (Next Advance). Each RINO sample tube (Next Advance) was supplemented with ~100 μl of stainless-steel beads (0.9–2.0 mm blend, Next Advance) and 500 μl of lysis buffer. Tissues were added immediately after excision and homogenized with Bullet Blender at 4 °C with two full 5-min cycles. The lysates were transferred to new Eppendorf LoBind tubes and sonicated for three cycles consisting of 5 s of active sonication at 30% amplitude, followed by 15 s on ice. Samples were then centrifuged for 5 min at 15,000*g* and the supernatant transferred to a new tube. Protein concentration was determined by bicinchoninic acid assay (Pierce) and one-dimensional SDS-PAGE gels were run followed by Coomassie blue staining as quality control for protein integrity and equal loading before proceeding to protein digestion.

### Brain protein digestion

For protein digestion (as described (18, 21, 70)), 100 μg of each sample was aliquoted, and volumes were normalized with additional lysis buffer. Samples were reduced with 1 mM dithiothreitol at room temperature for 30 min, followed by 5 mM iodoacetamide alkylation in the dark for another 30 min. Lysyl endopeptidase (Wako) at 1:100 (wt/wt) was added, and digestion was allowed to proceed overnight. Samples were then seven-fold diluted with 50 mM ammonium bicarbonate. Trypsin (Promega) was added at 1:50 (wt/wt), and digestion was carried out for another 16 h. The peptide solutions were acidified to a final concentration of 1% (vol/vol) formic acid (FA) and 0.1% (vol/vol) trifluoroacetic acid (TFA) and de-salted with a 30-mg HLB column (Oasis). Each HLB column was first rinsed with 1 ml of methanol, washed with 1 ml of 50% (vol/vol) acetonitrile (ACN) and equilibrated with 2× 1 ml of 0.1% (vol/vol) TFA. The samples were then loaded onto the column and washed with 2× 1 ml of 0.1% (vol/vol) TFA. Elution was performed with 2 volumes of 0.5 ml of 50% (vol/vol) ACN. An equal amount of peptide from each sample was aliquoted and pooled as the pooled global internal standard (GIS), which was split and labeled in each TMT batch as described below. The eluates were then dried to completeness using a SpeedVac.

### Brain Tandem Mass Tag (TMT) peptide labeling

Before TMT labeling, cases were randomized by covariates (age, sex, PMI, diagnosis, etc.), into the 26 total batches. Peptides from each individual case and the GIS pooled standard or bridging sample (at least one per batch) were labeled using the TMT 11-plex kit (ThermoFisher 90406). Labeling was performed as described (18, 70–72). In each batch, up to two TMT channels were used to label GIS standards, and the remaining TMT channels were reserved for individual samples after randomization. In brief, each sample (containing 100 μg of peptides) was re-suspended in 100 mM TEAB buffer (100 μl). The TMT labeling reagents (5 mg) were equilibrated to room temperature, and anhydrous ACN (256 μl) was added to each reagent channel. Each channel was gently vortexed for 5 min, and then 41 μl from each TMT channel was transferred to the peptide solutions and allowed to incubate for 1 h at room temperature. The reaction was quenched with 5% (vol/vol) hydroxylamine (8 μl) (Pierce). All channels were then combined and dried by SpeedVac (Labconco) to approximately 150 μl and diluted with 1 ml of 0.1% (vol/vol) TFA and then acidified to a final concentration of 1% (vol/vol) FA and 0.1% (vol/vol) TFA. Labeled peptides were de-salted with a 200-mg C18 Sep-Pak column (Waters). Each Sep-Pak column was activated with 3 ml of methanol, washed with 3 ml of 50% (vol/vol) ACN and equilibrated with 2× 3 ml of 0.1% TFA. The samples were then loaded, and each column was washed with 2× 3 ml of 0.1% (vol/vol) TFA, followed by 2 ml of 1% (vol/vol) FA. Elution was performed with 2 volumes of 1.5-ml 50% (vol/vol) ACN. The eluates were then dried to completeness using a SpeedVac.

### Brain high-pH offline fractionation

Fractionation was conducted as described (18, 71,73). Dried samples were re-suspended in high-pH loading buffer (0.07% v/v NH4OH, 0.045% v/v FA, 2% v/v ACN) and loaded onto an Agilent ZORBAX 300 Extend-C18 column (2.1 mm × 150 mm with 3.5 um beads). An Agilent 1100 HPLC system was used to carry out the fractionation. Solvent A consisted of 0.0175% (vol/vol) NH4OH, 0.01125% (vol/vol) FA, and 2% (vol/vol) ACN; solvent B consisted of 0.0175% (vol/vol) NH4OH, 0.01125% (vol/vol) FA, and 90% (vol/vol) ACN. The sample elution was performed over a 58.6 min gradient with a flow rate of 0.4 mL/min. The gradient consisted of 100% solvent A for 2 min, then 0% to 12% solvent B over 6 min, then 12% to 40 % over 28 min, then 40% to 44% over 4 min, then 44% to 60% over 5 min, and then held constant at 60% solvent B for 13.6 min. A total of 96 individual equal volume fractions were collected across the gradient and subsequently pooled by concatenation (73) into 24 fractions and dried to completeness using a SpeedVac.

### Brain liquid chromatography-tandem mass spectrometry

All fractions were resuspended in an equal volume of loading buffer (0.1% FA, 0.03% TFA, 1% ACN) and analyzed by LC–MS/MS essentially as described (18, 74) Peptide eluents were separated on a self-packed C18 (1.9 μm, Dr. Maisch) fused silica column (25 cm × 75 μM internal diameter, New Objective) by a Dionex UltiMate 3000 RSLCnano liquid chromatography system (Thermo Fisher Scientific) for the ROSMAP samples. Peptides were monitored on an Orbitrap Fusion mass spectrometer (Thermo Fisher Scientific). sample elution was performed over a 120-min gradient with flow rate of 300 nl min^−1^ with buffer B ranging from 1% to 50% (buffer A: 0.1% FA in water; buffer B: 0.1% FA in 80% ACN). The mass spectrometer was set to acquire in data-dependent mode using the top speed workflow with a cycle time of 3 s. Each cycle consisted of one full scan followed by as many MS/MS (MS2) scans that could fit within the time window. Full MS scans were collected at a resolution of 120,000 (400–1,400 *m*/*z* range, 4 × 10^5^ AGC, 50-ms maximum ion injection time). All HCD MS/MS spectra were acquired at a resolution of 60,000 (1.6 *m*/*z* isolation width, 35% collision energy, 5 × 10^4^ AGC target, 50-ms maximum ion time). Dynamic exclusion was set to exclude previously sequenced peaks for 20 s within a 10-ppm isolation window.

### Brain database searching and protein quantification

All raw MS data files (624 total RAW files generated across 26 batches) were analyzed in the Proteome Discover software suite (version 2.3, ThermoFisher) and MS/MS spectra were searched against the UniProtKB human proteome database (downloaded April 2015 with 90,411 total sequences). The Sequest HT search engine was used with the following parameters: fully tryptic specificity; maximum of two missed cleavages; minimum peptide length of 6; fixed modifications for TMT tags on lysine residues and peptide N-termini (+229.162932 Da) and carbamidomethylation of cysteine residues (+57.02146 Da); variable modification for oxidation of methionine residues (+15.99492 Da) and deamidation of asparagine and glutamine (+0.984 Da); precursor mass tolerance of 20 ppm; and fragment mass tolerance of 0.05 Da. Peptide spectral matches (PSMs) were filtered to a false discovery rate (FDR) of less than 1% using the Percolator node. Following spectral alignment, peptides were assembled into proteins and further filtered based on the combined probabilities of their constituent peptides to a final FDR of 1%. Multiconsensus was performed to achieve parsimony across individual batches. In cases of redundancy, shared peptides were assigned to the protein sequence in adherence with the principles of parsimony. Reporter ions were quantified from MS2 scans using integration tolerance of 20 ppm with the most confident centroid setting. Only unique and razor (i.e., parsimonious) peptides were considered for quantification.

### Brain data preprocessing

A total of 10,426 high confidence, master proteins were identified across all 26 TMTM batches, but only proteins quantified in >50% of samples were included in subsequent analyses (n=7,787 proteins). Log2 abundances were normalized as a ratio dividing by the central tendency of pooled standards (Global Internal Standards; GIS). As previously applied, batch correction was performed using a Tunable Approach for Median Polish of Ratio (https://github.com/edammer/TAMPOR; TAMPOR), an iterative median polish algorithm for removing technical variance across batch (18). Multidemensional scaling (MDS) plots were used to visualize batch contributions to variation before and after batch correction (**Fig. S1**). Network connectivity was used to remove outliers, that is samples that were greater than 3 standard deviations away from the mean as described (18). Finally, non-parametric bootstrap regression was performed to remove the potentially confounding covariates of age, sex and post-mortem interval (PMI). Each trait was subtracted times the median coefficient from 1000 iterations of fitting for each protein, while protecting for diagnosis (Control, AsymAD, AD).

### Brain Consensus Weighted Gene Correlation Network Analysis (cWGCNA)

We used the consensus Weighted Gene Correlation Network Analysis (cWGCNA; version 1.69) algorithm to generate a central network of co-expression modules from both brain regions (24, 25). The WGCNA::blockwiseConsensusModules function was run with soft threshold power at 7.0, deepsplit of 4, minimum module size of 30, merge cut height at 0.07, mean topological overlap matrix (TOM) denominator, using bicor correlation, signed network type, pamStage and pamRespectsDendro parameters both set to TRUE and a reassignment threshold of 0.05. This function calculates pair-wise biweight mid-correlations (bicor) between protein pairs. The resulting correlation matrix is then transformed into a signed adjacency matrix which is used to calculate a topological overlap matrix (TOM), representing expression similarity across samples for all proteins in the network. This approach uses hierarchical clustering analysis as 1 minus TOM and dynamic tree cutting lends to module identification. Following construction, module eigenprotein (ME) values were defined – representative abundance values for a module that also explain modular protein covariance. Pearson correlation between proteins and MEs was used as a module membership measure, defined as kME.

### Brain network preservation

We used the WGCNA::modulePreservation() function to assess the network module preservation of our current consensus network with recent large-scale TMT network from Brodmann area 9 (BA9) (18). Zsummary composite preservation scores were obtained using the consensus network as the test network and the previous BA9 TMT network as the reference network., with 500 permutations. Random seed was set to 1 for reproducibility, and the quickCor option was set to 0.

### Brain gene ontology (GO) and cell type marker enrichment analyses

To characterize differentially expressed proteins and co-expressed proteins based on GO annotation we used GO Elite (version 1.2.5) as previously described (18, 21, 75), with pruned output visualization using an in-house R script. Cell type enrichment was also investigated as previously published (18, 21,75). An in-house marker list combined previously published cell type marker lists from Sharma et al. (76) and Zhang t al. (77) were used for the cell type marker enrichment analysis for each of the five cell types assessed (neuron, astrocyte, microglia, oligodendrocyte and endothelial; **Supplemental Table 7**). If, after the lists from Sharma et al. and Zhang et al. were merged, gene symbol was assigned to two cell types, we defaulted to the cell type defined by the Zhang et al. list such that each gene symbol was affiliated with only one cell type. The gene symbols in the list were processed through MyGene to ensure updated nomenclature and then converted human symbols using homology lookup. Fisher’s exact tests were performed using the human cell type marker lists to determine cell type enrichment and were corrected by the Benjamini-Hochberg procedure (**Supplemental Table 8**).

### Proteome Wide Association Study (PWAS) results module enrichment analysis

Proteins (*n* = 8,356) tested in the PWAS study by Yu et al. (51) for correlation to cognitive resilience (or decline, when negatively correlated) were split into lists of unique gene symbols representing protein gene products positively correlated (*n* = 645) and negatively correlated (*n* = 575) to cognitive resilience, and then these lists with corresponding *P* values were separately checked for enrichment in consensus TMT network modules using a permutation-based test (10,000 permutations) implemented in R with exact *P* values for the permutation tests calculated using the permp function of the statmod package. Module-specific mean *P* values for risk enrichment were determined as a *Z* score, specifically as the difference in mean *P* value of gene product proteins hitting a module at the level of gene symbol minus the mean *P* value of genes hit in the 10,000 random replacement permutations, divided by the standard deviation of *P* value means also determined in the random permutations (**Supplemental Tables 9 and 10**).

### Thioflavin T aggregation assay

The effect of NRN1 on amyloid beta 1-42 (Aβ_42_) aggregation was measured by *in vitro* thioflavin T (ThT) fluorescence assay essentially as previously described (67). Recombinant human Aβ_42_ (20 μg/mL equivalent to 5 μM) from rPeptide (# A-1170-1) was incubated in 1× Tris-buffered Saline (TBS; 150 mM NaCl, 50 mM Tris-HCl, pH 7.6), and 20 μM ThT in the presence or absence of purified recombinant NRN1 (5 μg/mL or 263 nM; Abcam, ab69755) protein. The assay was conducted in 100 μL reaction volumes in quadruplicates using chilled 96 well black clear bottom plates (Corning, #3904). Fluorescence was captured at 420 Ex, 480 Em for 20 hours at 15 min intervals at 37°C using Synergy H1 (Biotek) microplate reader. ThT alone was measured and subtracted as background fluorescence. Fluorescence intensities were graphed using GraphPad prism.

### SDS-PAGE and immunoblot analyses

For human brain homogenates, 10 μg of protein from each sample was mixed with Laemmli sample buffer (Bio-rad) and β-mercaptoethanol, boiled at ~95° C for 10 minutes, spun briefly to collect the volume and loaded into Bolt 4-12% Bis-Tris gels (Invitrogen) and electrophoresed at 160 V for ~30 minutes. Gels were then stained with Coomassie Blue for protein banding visualization.

For products of the ThT aggregation assay, Aβ_42_ fibrils were precipitated by centrifugation at 10,000 × g. The pellet was resuspended in 50 μL 8M urea buffer (8 M urea, 100 mm NaHPO4, pH 8.5) and boiled in Laemmli sample buffer (BioRad, 161-0737) at 98°C for 5 min. Proteins were resolved on Bolt 4-12% Bis-Tris gels (Thermo Fisher Scientific, NW04120BOX) followed by transfer to nitrocellulose membrane using iBlot 2 dry blotting system (ThermoFisher Scientific, IB21001). Membranes were incubated with StartingBlock buffer (ThermoFisher, 37543) for 30 min followed by overnight incubation at 4° in primary antibodies, Aβ (Novus, NBP11 -97929) and NRN1 (Abcam, ab64186). Membranes were washed with 1×Tris-buffered saline containing 0.1% Tween 20 (TBS-T) and incubated with fluorophore-conjugated secondary antibodies (AlexaFluor-680 or AlexaFluor-800) for 1 h at room temperature. Membranes were subsequently washed three times with TBS-Tween and images were captured using an Odyssey Infrared Imaging System (LI-COR Biosciences).

### Silver staining

Aβ_42_ fibrils prepared in the ThT assay above were precipitated by centrifugation at 10,000 × g. The pellet was resuspended in 50 μL 8M urea buffer (8 M urea, 100 mm NaHPO4, pH 8.5) and 10 μL of fibrils were boiled in Laemmli sample buffer (BioRad, 161-0737) at 98°C for 5 min. Fibrils were run on Bolt 4-12% Bis-Tris gels (Thermo Fisher Scientific, NW04120BOX) and stained using a silver staining kit (Pierce, 24612) following manufacturers protocols. Briefly, the was rinsed twice in ultrapure water for 5 minutes followed by fixation in 30% ethanol, 10% acetic acid in water. The gel was washed in 10% Ethanol and water. The gel was then incubated in silver stain and developer solutions. Staining was quenched using 5% acetic acid and images were captured using a scanner.

### Primary rat hippocampal culture

Primary rat hippocampal cultures were generated from E18 Sprague-Dawley rat embryos as previously described (Swanger, Mattheyses et al. 2015, Henderson, Greathouse et al. 2019) Briefly, cell culture plates were coated overnight with 1 mg/mL poly-L-lysine (Sigma-Aldrich, catalog no. P2636-100MG) and rinsed with diH20. Neurons were cultured at a density of 4 × 10^5^ cells per 18-mm glass coverslip in 12-well culture plates (Fisher Scientific, catalog no. 353043). Neurons were cultured in Neurobasal medium (Fisher Scientific, catalog no. 21103-049) supplemented with B27 (Fisher Scientific, catalog no. 17504-044), conditioned by separate cultures of primary rat astrocytes and glia, in a humidified CO2 (5%) incubator at 37°C. Neurons were treated at DIV 4 with 5μM cytosine β-D-arabinofuranoside hydrochloride (Sigma-Aldrich, catalog no. C6645) to eliminate the presence of native astrocytes and glia on the glass coverslips. Medium was changed every three to four days with new glia-conditioned Neurobasal medium for proper culture maintenance. At DIV 12, neurons were transfected with plasmid using Lipofectamine 2000 (Invitrogen, catalog no. 11-668-019) according to the manufacturer’s instructions. At DIV 14, primary hippocampal neurons were dosed with either DMSO, 500nM Aβ_42_, 150 ng/mL recombinant neuritin (NRN1), or a combination of 500nM Aβ_42_ plus 150 ng/mL NRN1 for 6 hours. 6 hours was chosen based on past studies demonstrating that Aβ_42_-induced spine loss in cultured neurons plateaus at approximately 6 hours post exposure(30, 32).

### Static widefield microscopy

On DIV 14, neurons were fixed with room temperature 2% paraformaldehyde (PFA) in 0.1M phosphate-buffered saline (PBS), washed two times with 1X PBS, and coverslips were mounted on microscope slides (Fisher Scientific, catalog no. 12-550-15) using Vectashield mounting media (Vector Labs, catalog no. H1000). A blinded experimenter performed all microscopy. Images were captured on a Nikon (Tokyo, Japan) Eclipse Ni upright microscope, using a Nikon Intensilight and Photometrics Coolsnap HQ2 camera to image Lifeact-GFP. Previous studies demonstrated that Lifeact-expressing neurons display normal, physiological actin dynamics and dendritic spine morphology (78, 79). Images were captured with Nikon Elements 4.20.02 image capture software using 60X oil-immersion objective (Nikon Plan Apo, N.A. 1.40). Z-series images were acquired at 0.10μm increments through the entire visible dendrite. Dendrites were selected for imaging by using the following criteria: 1) minimum of 25μm from the soma; 2) no overlap with other branches; 3) must be a secondary dendritic branch. Prior to analysis, capture images were deconvolved using Huygens Deconvolution System (16.05, Scientific Volume Imaging, the Netherlands) with the following settings: CMLE; maximum iterations: 50; signal to noise ratio: 40; quality: 0.1. Deconvolved images were saved in .tif formation.

### Dendritic spine morphometry analysis

Image analysis was performed with Neurolucida 360 (2.70.1, MBD Biosciences, Williston, Vermont) based on previously described methods(32). Dendritic spine reconstruction was performed automatically using a voxel-clustering algorithm and the following parameters: outer range: 10.0μm; minimum height: 0.5μm; detector sensitivity 100%; minimum count: 8 voxels. Next, the experimenter manually verified that the classifier correctly identified all protrusions. When necessary, the experimenter added any protrusions semi-automatically by increasing detector sensitivity. Each dendritic protrusion was automatically classified as a dendritic filopodium, thin spine, stubby spine, or mushroom spine based on previously described morphological measurements(80). Reconstructions were collected in Neurolucida Explorer (2.70.1, MBF Biosciences, Williston, Vermont) for branched structure analysis, and then exported to Microsoft Excel (Redmond, WA). Spine density was calculated as the number of spines per 10μm of dendrite length.

### Multi-electrode array recording and analysis

Single neuron electrophysiological activity was recorded using a Maestro Edge multiwell microelectrode array and Impedance system (Axion Biosystems). 24 hours prior to multielectrode array (MEA) culturing, each well of a 6-well plate (Axion Biosystems, catalog no. M384-tMEA-6W-5) was coated with 1 mg/mL Poly-L-lysine (Sigma, catalog no. P2636-100MG). The next day, wells were washed with diH_2_O. E18 rat primary hippocampal neurons were harvested as described above and plated in a 6-well MEA at a density of 4 x 10^5^ cells per well. Each MEA well contained 64 extracellular recording electrodes. Neurons were cultured DIV 0 to DIV 4 in Neurocult™ Neuronal Plating Medium (Stemcell Technologies, catalog no. 05713) with SM1 neuronal supplement (Stemcell Technologies, catalog no. 05711). At DIV 4, media was changed to BrainPhys™ Neuronal Medium (Stemcell Technologies, catalog no. 05790) with SM1 neuronal supplement. At DIV 14, a 5-min MEA prerecording was performed followed by application of DMSO, 500nM Aβ_42_, 150 ng/mL NRN1, or 150 ng/mL NRN1 and 500nM Aβ_42_. After 6 hours, a follow-up 5-min MEA recording was performed to determine effects on neuronal firing. All recordings were performed while connected to a temperature-controlled heater plate (37°C) with 5% CO_2_. All data were filtered using 0.1-Hz (high pass) and 5-kHz (low pass) Butterworth filters. Action potential thresholds were set manually for each electrode (typically > 6 standard deviations from the mean signal). Sorting of distinct waveforms corresponding to multiple units on one electrode channel were completed in Offline Sorter (v. 4.0, Plexon). Further analysis of firing rate was performed in NeuroExplorer (v. 5.0, Plexon). Mean firing frequency was calculated spikes/second and log_10_ transformed.

### Cortical rat neuronal culture, lysis and proteolytic digestion

Primary rat cortical neurons were generated from E18 Sprague-Dawley rat embryos with minor modifications (Swanger, Mattheyses et al. 2015, Henderson, Greathouse et al. 2019). Neurons were cultured at a density of 4×10^5^ cells per well in 12-well culture plates (Fisher Scientific, catalog no. 353043). Neurons were cultured in Neurobasal medium (Fisher Scientific, catalog no. 21103-049) supplemented with B27 (Fisher Scientific, catalog no.17504-044). Culture maintenance included a half media change every 2-3 days. At DIV 14, neurons were either treated with 150 ng/mL recombinant NRN1 protein (Abcam, ab69755) or vehicle treated with diH2O for 6 hours. NRN1 concentration was chosen based on published data that identified a plateau in exogenous NRN1 induced effects on transient potassium currents at 150 ng/mL (81). After 6 hours neurons were washed 2× with 1 mL 1X phosphate-buffered saline (PBS). To harvest cells, 1 mL 1X PBS + protease inhibitor (Fisher Scientific, catalog no. 78426) was added and cells were centrifuged for 2300rpm for 5 minutes at 4°C. Cell pellets were lysed in 200uL 8M urea buffer and HALT protease and phosphatase inhibitor cocktail (1× final concentration). Lysates were sonicated with a probe sonicator 3 times for 10 s with 10 s intervals at 30% amplitude and cleared of cellular debris by centrifugation in a tabletop centrifuge at 18,000 rcf for 3 minutes at 4° C. Protein concentration was determined by BCA assay and one-dimensional SDS-PAGE gels were run followed by Coomassie blue staining as quality control for protein integrity and equal loading before proceeding to protein digestion. Protein homogenates (50 μg) were diluted with 50 mM NH4HCO3 to a final concentration of less than 2 M urea and then treated with 1 mM dithiothreitol (DTT) at 25°C for 30 minutes, followed by 5 mM iodoacetimide (IAA) at 25°C for 30 minutes in the dark. Protein was digested with 1:100 (w/w) lysyl endopeptidase (Wako) at 25°C for 2 hours and further digested overnight with 1:50 (w/w) trypsin (Pierce) at 25°C. Resulting peptides were desalted with a Sep-Pak C18 column (Waters) and dried under vacuum.

### Rat neuron TMT labeling

Peptides from each individual cell line in the study and a global pooled reference internal standard (GIS) were labeled using the TMTpro 16-plex kit (ThermoFisher Cat#A44520 Lot#VH311511). Labeling was performed essentially as previously described (19, 82). Briefly, each sample (containing 100 μg of peptides) was re-suspended in 100 mM TEAB buffer (100 μL). The TMT labeling reagents were equilibrated to room temperature, and anhydrous ACN (256 μL) was added to each reagent channel. Each channel was gently vortexed for 5 min, and then 41 μL from each TMT channel was transferred to the peptide solutions and allowed to incubate for 1 h at room temperature. The reaction was quenched with 5% (vol/vol) hydroxylamine (8 μl) (Pierce). All 16 channels were then combined and dried by SpeedVac (LabConco) to approximately 150 μL and diluted with 1 mL of 0.1% (vol/vol) TFA, then acidified to a final concentration of 1% (vol/vol) FA and 0.1% (vol/vol) TFA. Peptides were desalted with a 200 mg C18 Sep-Pak column (Waters). Each Sep-Pak column was activated with 3 mL of methanol, washed with 3 mL of 50% (vol/vol) ACN, and equilibrated with 2×3mL of 0.1% TFA. The samples were then loaded, washed with 2×3 mL 0.1% (vol/vol) TFA and 2 mL of 1% (vol/vol) FA. Elution was performed with 2 volumes of 1.5 mL 50% (vol/vol) ACN. The eluates were then dried to completeness. High pH fractionation was performed next as described for human samples.

### Rat neuron LC-MS/MS

All samples were analyzed with a Dionex Ultimate 3000 RSLCnano in capillary flow mode. The analytical column was a 300 μm × 150 mm ID Waters CSH with 1.7 μm beads. Mass spectrometry was performed with a high-field asymmetric waveform ion mobility spectrometry (FAIMS) Pro equipped Orbitrap Eclipse (Thermo) in positive ion mode using data-dependent acquisition with 1.5 second top speed cycles for each FAIMS compensation voltage (CV). Each cycle consisted of one full MS scan followed by as many MS/MS events that could fit within the given 1.5 second cycle time limit. MS scans were collected at a resolution of 120,000 (410-1600 m/z range, 4×10^5^ AGC, 50 ms maximum ion injection time, FAIMS CV of −45 and −65). All higher energy collision-induced dissociation (HCD) MS/MS spectra were acquired at a resolution of 30,000 (0.7 m/z isolation width, 35% collision energy, 1.25×10^5^ AGC target, 54 ms maximum ion time, TurboTMT on). Dynamic exclusion was set to exclude previously sequenced peaks for 20 seconds within a 10-ppm isolation window. All raw files are loaded onto synapse folder https://www.synapse.org/ADresilienceRat.

### Rat neuron data search and protein quantification

All raw files (n=96) were analyzed using the Proteome Discoverer Suite (version 2.4) Thermo Scientific). MS/MS spectra were searched against the UniProtKB rat proteome database (downloaded April 2015 with 29370 total sequences). The Sequest HT search engine was used with the following parameters: fully tryptic specificity; maximum of two missed cleavages; minimum peptide length of 6; fixed modifications for TMT tags on lysine residues and peptide N-termini (+304.207 Da) and carbamidomethylation of cysteine residues (+57.02146 Da); variable modifications for oxidation of methionine residues (+15.99492 Da), deamidation of asparagine and glutamine (+0.984 Da) and phosphorylation of serine, threonine and tyrosine (+79.966); precursor mass tolerance of 10 ppm; and fragment mass tolerance of 0.05 Da. The Percolator node was used to filter peptide spectral matches (PSMs) to a false discovery rate (FDR) of less than 1%. Following spectral assignment, peptides were assembled into proteins and were further filtered based on the combined probabilities of their constituent peptides to a final FDR of 1%. A Multi-consensus was performed to group proteins identified across the individual batches. In cases of redundancy, shared peptides were assigned to the protein sequence in adherence with the principles of parsimony. A total of 125869 peptides mapping to 9799 protein groups. Reporter ions were quantified from MS2 scans using an integration tolerance of 20 ppm with the most confident centroid setting. Only unique and razor (i.e., parsimonious) peptides were considered for quantification. TMT channels 129C, 130N and 130C correspond to NRN1 treated samples and channels 132C, 133N, 133C and 134N correspond to vehicle treated samples which were used for the presented results (**Supplemental Table 13**; https://www.synapse.org/ADresilienceRat).

### Rat neuronal proteome overlap with human consensus modules

Human consensus module (39 modules) protein members were converted to rat symbols using the biomaRt package and overlap of rat neuronal proteins was determined for each module. A one-tailed Fisher exact test looking for significant overrepresentation or overlap was employed, and *P* values were corrected for multiple testing using the Benjamini–Hochberg method. R functions fisher.test() and p.adjust() were used to obtain the above statistics (**Supplemental Table 15**).

### Additional statistical analyses

All proteomic statistical analyses were performed in R (version 4.0.3). Box plots represent the median and 25^th^ and 75^th^ percentile extremes; thus the hinges of a box represent the interquartile range of the two middle quartiles of data within a group. Error bars extents are defined by the farthest data points up to 1.5 times the interquartile range away from the box hinges. Correlations were performed using biweight midcorrelation function from the WGCNA package. Group comparisons in human brain samples were performed with one-way ANOVA with Holm post hoc correction of all comparisons. Differential expression between NRN1 and vehicle treated neurons was determined by student’s t-test (**Supplemental Table 14**). Differential expression displayed as volcano plots were generated using the ggplot2 package. Go annotation for rat neuron proteins was performed as described for human samples. P values were adjusted for multiple comparisons by FDR correction where indicated.

All analyses from dendritic spine morphometric and MEA results were conducted with Prism 9.0 (GraphPad Software, La Jolla, CA). Data are presented as mean + SEM, and all graph error bars represent SEM. All statistical tests were two tailed with threshold for statistical significance set at 0.05. Statistical comparisons on spine densities and morphologies are oneway ANOVA with Tukey’s comparison’s test. Statistical comparisons on mean firing rate are unpaired Student’s *t* test.

## Acknowledgements

This study was supported by the following Nation Institutes of Health funding mechanisms: NINDS T32 NS061788 (DAP), NIA AG054719 (JHH), NIA AG063755, and NIA AG068024, R01AG061800 (J.H.H and N.T.S) and U01AG061357 (N.T.S).

## Author contributions

C.H., D.A.P., N.T.S. and J.H.H. designed experiments. C.H., D.A.P., M.A., and D.M.D carried out experiments. C.H and D.A.P analyzed data. N.T.S, J.H.H., and E.B.D. provided advice on interpretation of data. C.H. and D.A.P. wrote the manuscript with input from all co-authors. D.A.B provided human tissue samples. All authors approved the final manuscript.

## Supplemental Figure Legends

**Supplemental Figure S1.**
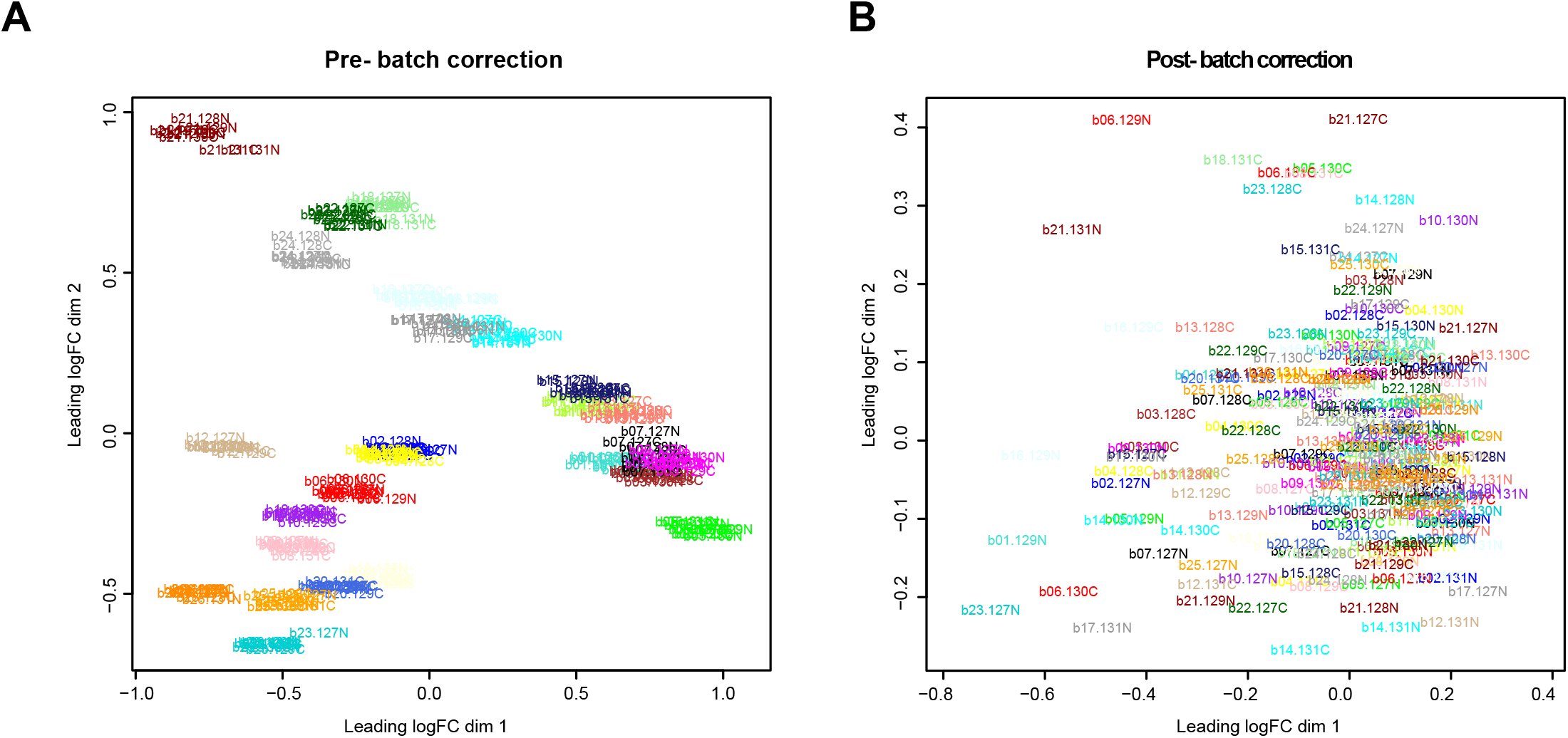
TMT batch correction across BA6 and BA37. A median polish batch correction approach was implemented to remove technical batch variance across the 26 TMT 11-plex batches. (**A**) Multidimensional scaling (MDS) plots visualize original log_2_ transformed protein abundances, normalized to the pooled global internal standards (GIS). (**B**) MDS of batch-corrected normalized log_2_ abundance after 175 iterations. Samples are color-coded by batch.

**Supplemental Figure S2.**
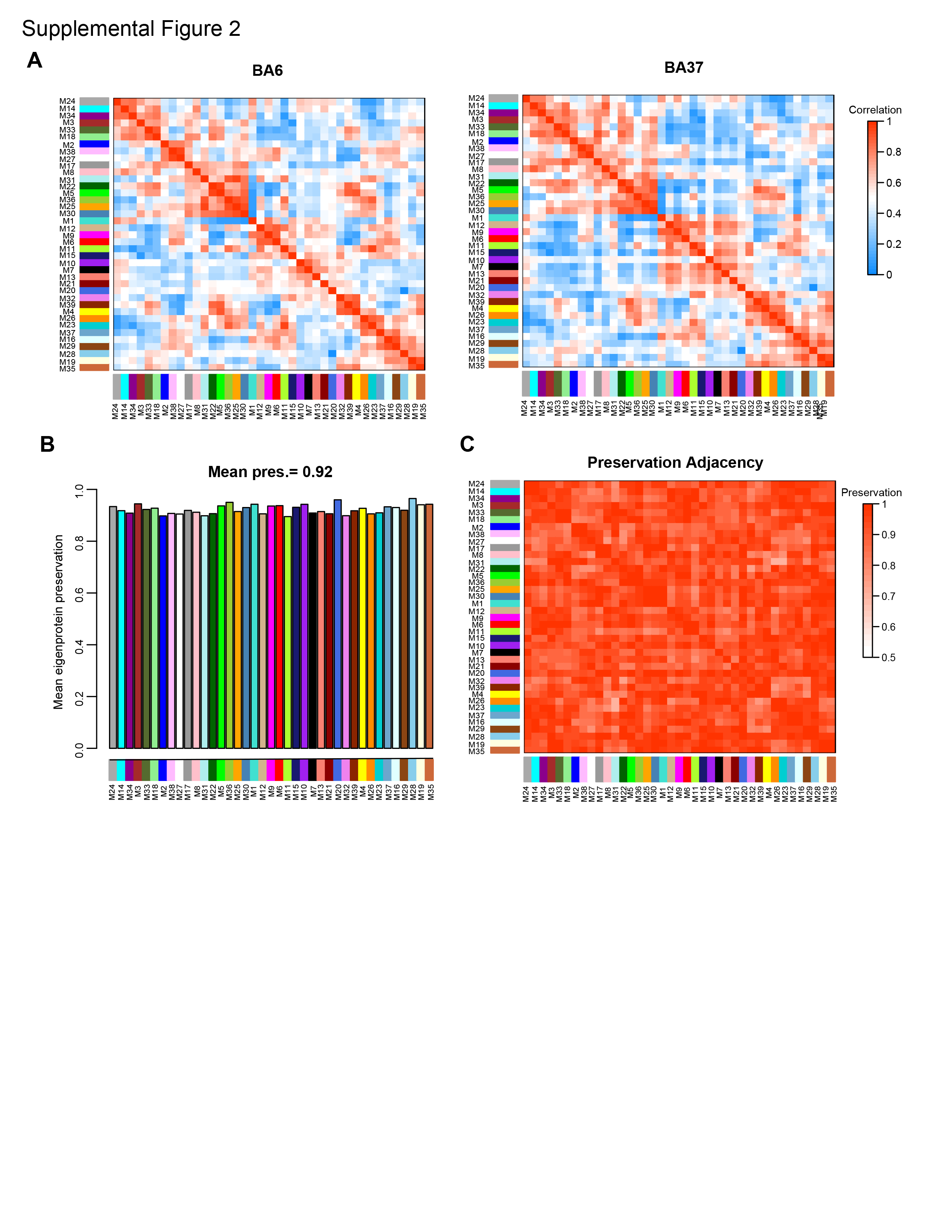
Consensus modules are highly preserved in BA6 and BA37. (**A**) Modules eigenproteins were correlated to visualize inter-module relationships in BA6 and BA37, respectively. Heat blocks along the diagonal could be observed similarly in both brain regions. (**B**) Mean preservation relationship for each eigenprotein was calculated for the consensus network, with mean preservation of 0.92 indicating very high preservation. (**C**) Preservation adjacency of the consensus network, visualized as a heatmap, further support that most relationships in the network across both brain regions are highly preserved and biologically meaningful.

**Supplemental Figure S3.**
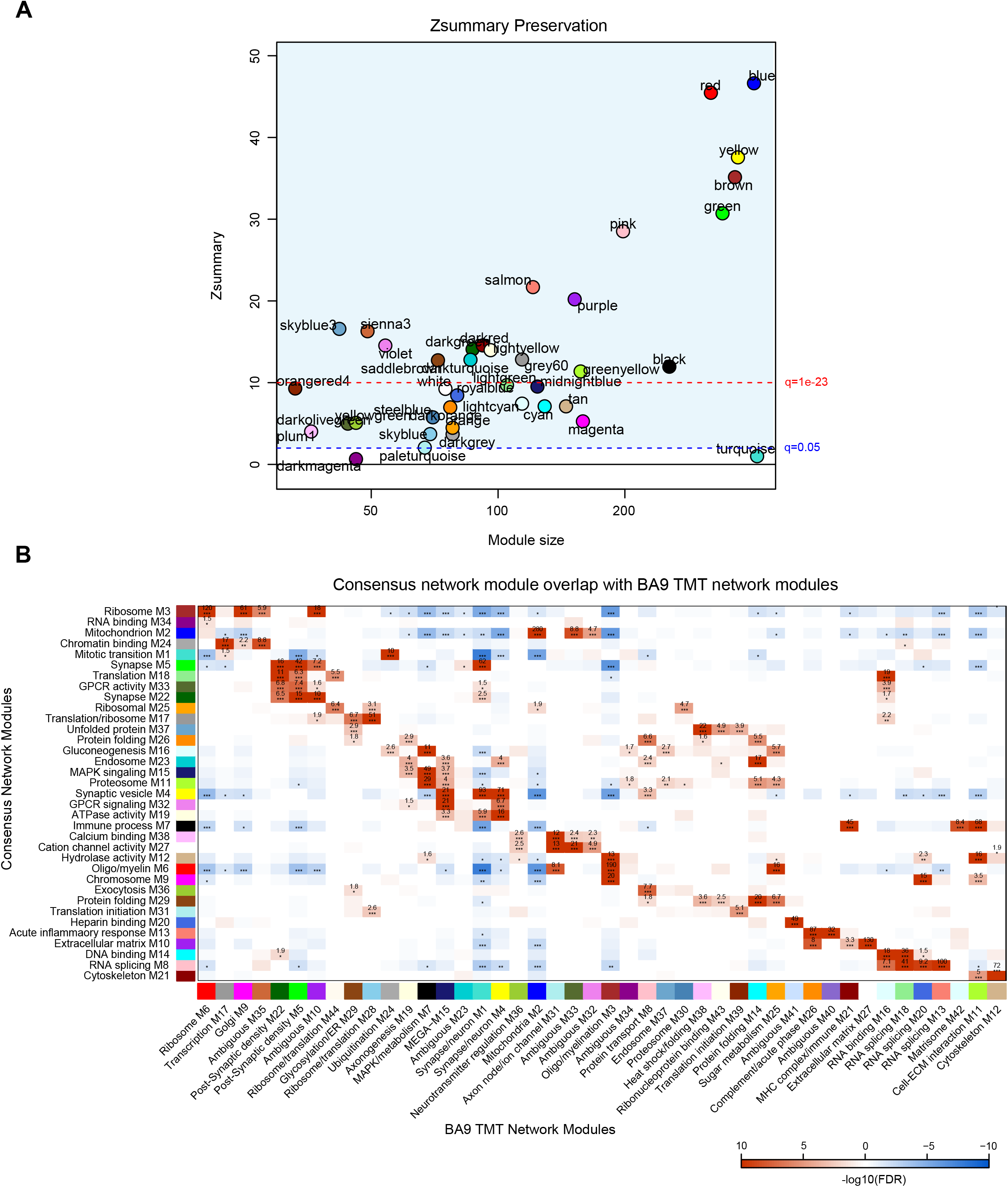
Consensus network preservation. (**A**) Zsummary indicates nearly all consensus modules preserve with previous BA9 TMT network modules reported in Johnson et al. Module Zsummary greater than or equal to 1.96 (q=0.05, dashed blue line) are considered preserved and modules with Zsummary of 10 or higher (q=1e-23, dashed red line) are considered highly preserved. (**B**) Overrepresentation analysis of consensus modules members with previous BA9 TMT module members. -log_10_ FDR corrected overlap values are shown. The heatmap threshold is at a 10% FDR (0.1).

**Supplemental Figure S4.**
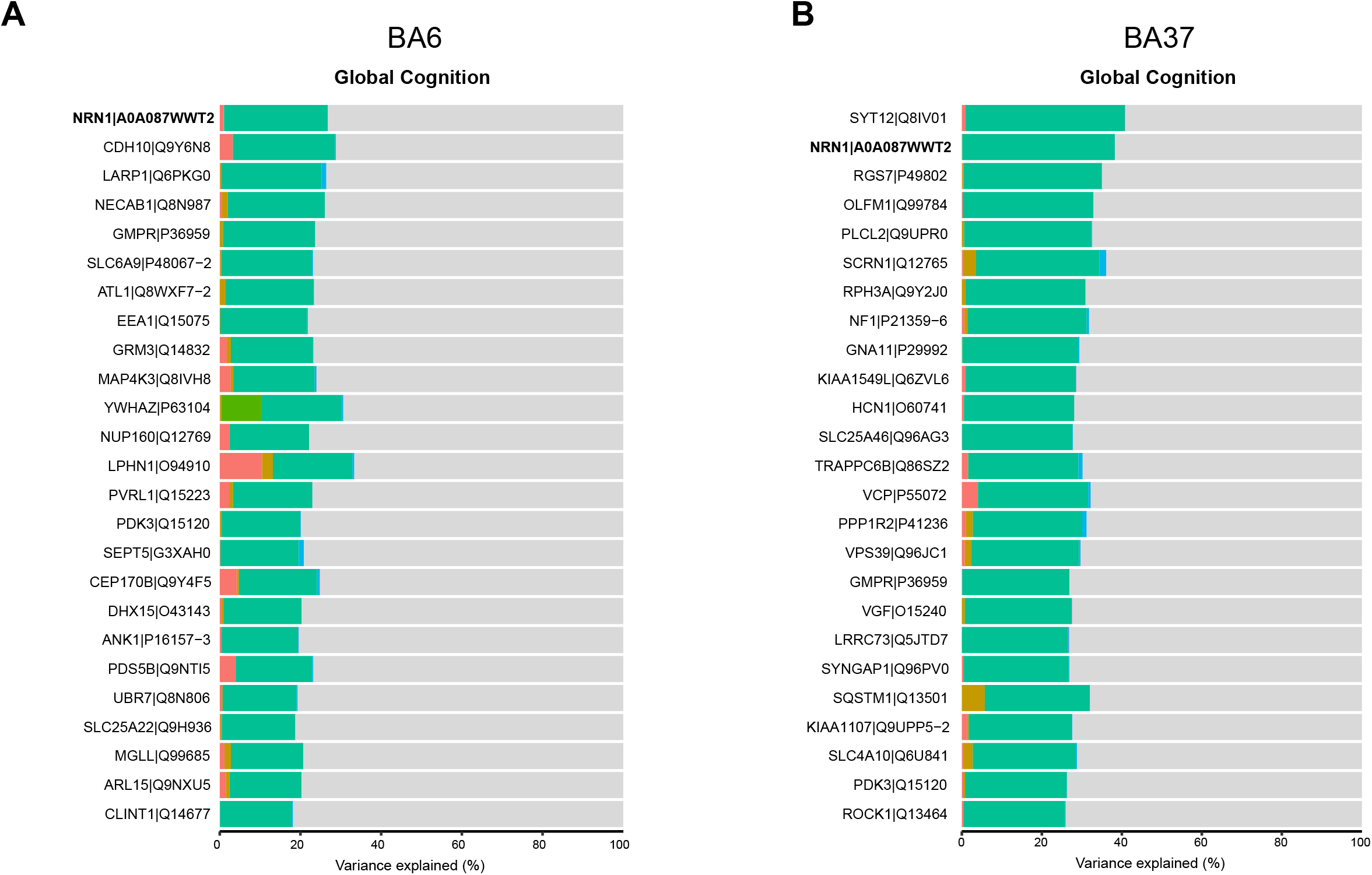
Percent variance in protein expression explained by global cognition. A linear mixed model approach was implemented to estimate the percent variance explained by proteins in relationship to diagnosis, CERAD score, Braak score and global cognition across region, respectively. The rank order in the percent variation in protein expression explained by global cognition (top 20 proteins) were plotted for BA6 (**A**) and BA37 (**B**).

**Supplemental Figure S5.**
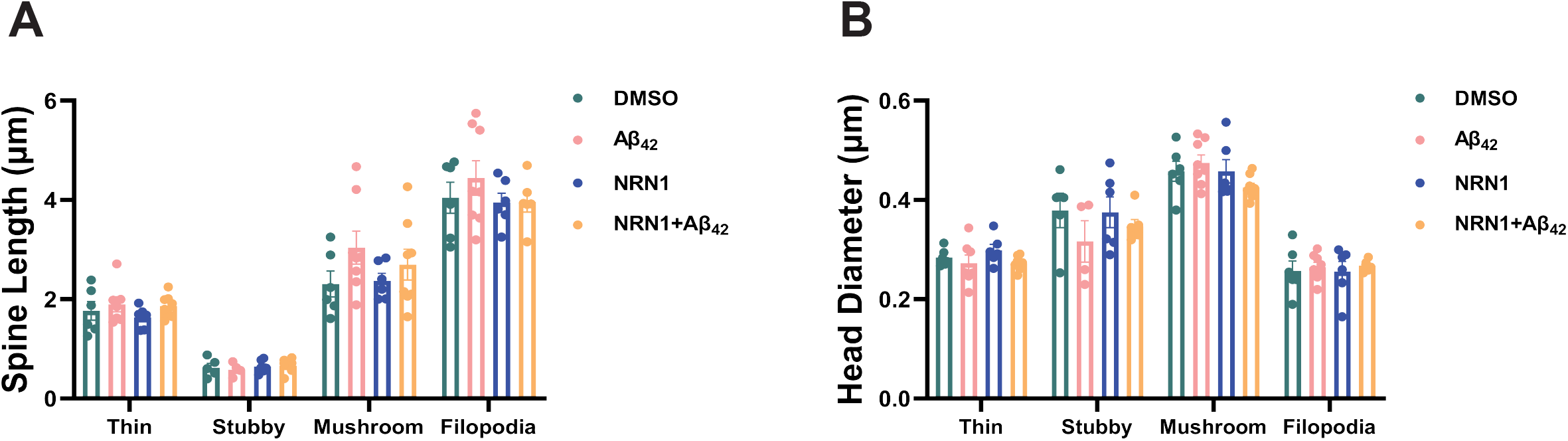
Analysis of dendritic spine length and head diameter among thin, stubby, and mushroom spines, and filopodia. (**A**) Mean dendritic spine length of thin, stubby, or mushroom spines, and filopodia. Data are means + SEM. (**B**) Mean dendritic spine head diameter of thin, stubby, or mushroom spines, and filopodia. Data are means + SEM.

**Supplemental Figure S6.**
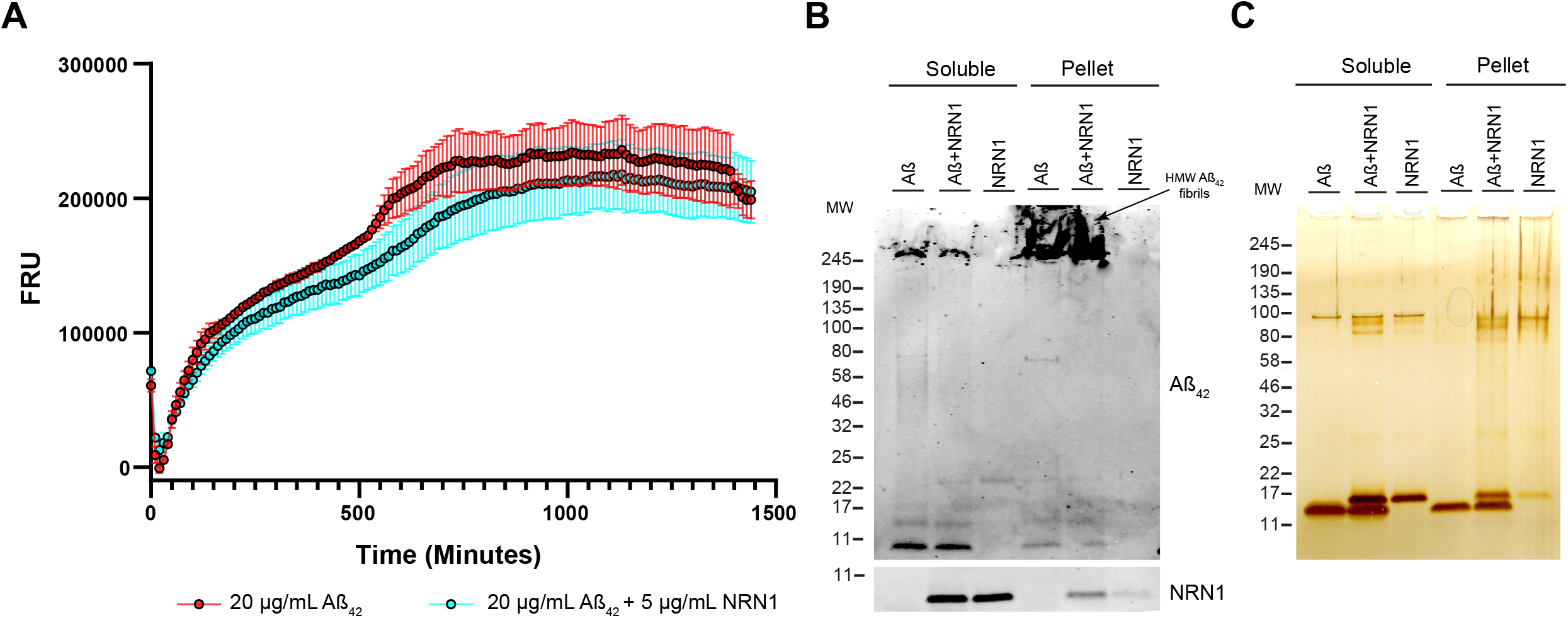
Aggregation of Aβ in the presence or absence of NRN1. (**A**) Fibrillation curves of 20 μg/mL Aβ_42_ alone and 20 μg/mL Aβ_42_ + 5 μg/mL NRN1, thioflavin T (ThT) alone was recorded and subtracted as background. Relative fluorescent units (RFU) were recorded every 15 minutes for 20 hours. Points are quadruplicate means ± SEM. (B) Western blot of soluble and pelleted fractions of assay products probed for Aβ_42_ and NRN1. High molecular weight (HMW) fibrils are observed at the top of the gel. (C) Silver stain of soluble and pelleted fractions of assay products.

**Supplemental Figure S7.**
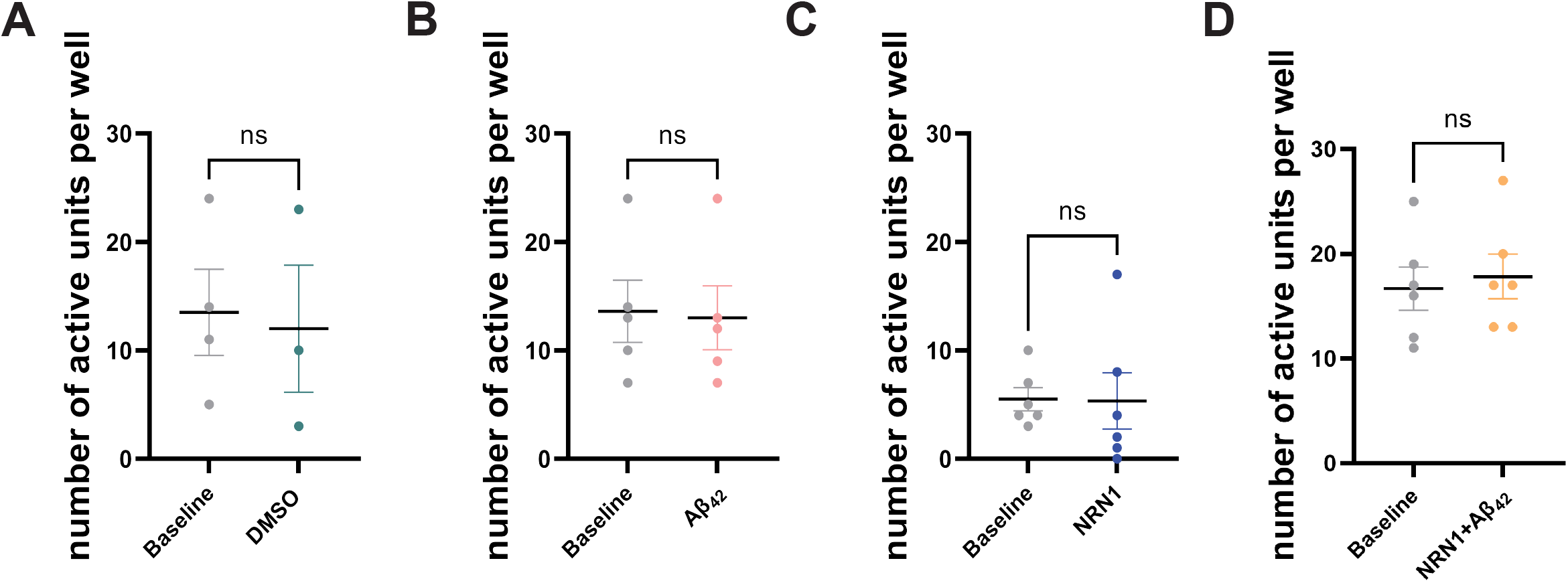
Total number of active neurons per microelectrode array. (**A**) The total number of active neurons per well at DIV 14 in hippocampal neurons treated with DMSO, compared to baseline (n = 3-4 wells with 64 electrodes/well, unpaired Student’s t test; p = 0.8339). Data are means + SEM. (**B**) The total number of active neurons per well at DIV 14 in hippocampal neurons treated with 500nM Aβ_42_, compared to baseline (n = 5 wells with 64 electrodes/well, unpaired Student’s t test; p = 0.8878). Data are means + SEM. (**C**) The total number of active neurons per well at DIV 14 in hippocampal neurons treated with 150 ng/mL NRN1, compared to baseline (n = 6 wells with 64 electrodes/well, unpaired Student’s t test; p = 0.9539). Data are means + SEM. (**D**) The total number of active neurons per well at DIV 14 in hippocampal neurons treated with 150 ng/mL NRN1 and 500nM Aβ_42_, compared to baseline (n = 6 wells with 64 electrodes/well, unpaired Student’s t test; p = 0.7035). Data are means + SEM.

